# Climate change impacts dryland multifunctionality: A cascade of precipitation variability to species replacement in multitrophic microbiota

**DOI:** 10.1101/2023.12.19.572331

**Authors:** Xiaoyu Guo, Hua Li, Zuowen Wang, Haijian Yang, Chunxiang Hu, Lirong Song, Weibo Wang

## Abstract

In the Anthropocene, the pervasive impacts of climate change on arid ecosystems are increasingly evident, resulting in significant alterations in both biodiversity and functionality. Although soil microbes are anticipated to be sensitive to changes in precipitation regimes, the mechanisms through which multifaceted precipitation changes impair the delivery of multiple functions by multitrophic microbiota in soils remain largely unexplored. In the drylands of Northwestern China, we examined the direct effects of historical precipitation regimes on the soil multifunctionality of biocrusts, a model system, and their indirect impacts mediated by microbial communities. Among abiotic predictors, precipitation variability, rather than mean annual precipitation, emerged as the primary driver of multifunctionality, leading to a convergence of less functionally beneficial species. By utilizing a biodiversity-ecosystem function relationship framework, our results underscore the crucial roles of *α*- and *β*-diversity of heterotrophic bacteria, fungi, and phototrophic cyanobacteria in maintaining soil functionality. However, the underlying mechanisms are distinct. We found that the local richness of heterotrophs and phylogenetic dissimilarity of photoautotrophs exert positive influences on multifunctionality, while species replacement, a component of *β*-diversity, primarily enhances the variance in soil multifunctionality. Importantly, the findings illuminate the cascading effect of precipitation variability on dryland ecosystems, amplified by microbial diversity, potentially triggering a silent collapse of vulnerable arid habitats. Our study highlights the importance of considering indirect climate impacts via soil microbiota, contributing to a deeper understanding of the real-world consequences of climate change on drylands, and offering valuable insights for the management of biodiversity theory-inspired ecosystem restoration.

## Introduction

Climate change has instigated significant shifts in precipitation regimes on both global and regional scales in recent decades (Lenderink & Fowler, 2017; Ombadi, 2023). These alterations directly affect soil moisture levels and availability (Haiyan & Haimei, 2021), potentially triggering a cascade of ecological consequences, including changes in biodiversity and ecosystem functions (Austin et al., 2004; J. W. Zhang et al., 2023). The effects are anticipated to be particularly pronounced in water-limited environments such as arid and semi-arid regions, where ecosystems, due to long-term low precipitation inputs, exhibit heightened sensitivity to precipitation changes (Maurer, Hallmark, Brown, Sala, & Collins, 2020). Furthermore, historical precipitation patterns may exert legacy effects on ecological properties, thereby intensifying their impact on the biotic community and corresponding functionality (Canarini et al., 2021; S. Evans, Allison, & Hawkes, 2022). Despite recent studies beginning to document dryland biome responses to precipitation shifts (Holguin, Collins, & McLaren, 2022; Huang, Li, & Su, 2015; Nielsen & Ball, 2015; Wu et al., 2022), there remains an urgent need to explore the response and functional impact framework of soil microbiota in these regions.

Global circulation models (GCMs) predict an escalation of extreme events, such as variable precipitation and severe, prolonged droughts, in drylands worldwide (J. B. Bradford, Schlaepfer, Lauenroth, & Palmquist, 2020; IPCC, 2021). These historical precipitation changes can lead to landscape-scale homogenization or heterogeneity of precipitation regimes among arid and semi-arid regions (Shang, Xu, Zhao, & Tijjani, 2019). Given the mega-diverse and crucial roles of soil microorganisms in the stability and resilience of terrestrial ecosystems (Wagg et al., 2021), recent research has shifted focus toward their responses to precipitation alterations (Cregger, Schadt, McDowell, Pockman, & Classen, 2012; S. E. Evans & Wallenstein, 2014; Xi & Bloor, 2016). For example, increased precipitation has been found to elevate the fungi-to-bacteria ratio, potentially influencing the resistance of specific ecosystem functions (Bell et al., 2014; Bi, Zhang, Liang, Yang, & Ma, 2012). In contrast, decreased precipitation has been associated with negative impacts on microbial diversity, network complexity, and soil multifunctionality (X. Wang et al., 2023). However, beyond the amount of precipitation (*e.g.*, mean annual precipitation, MAP), the response of soil microbial communities to altered precipitation variability remains largely unknown (Chen, Huo, Zhang, Guo, & Liang, 2023), despite predictions of its universal increase in the future (W. Zhang et al., 2021). Notably, the delivery pattern of precipitation can have significant consequences, particularly in vulnerable habitats (Austin et al., 2004; Nielsen & Ball, 2015). Therefore, a comprehensive evaluation of the impacts of multiple facets of precipitation regime changes on dryland microbiota at different trophic levels is increasingly essential.

Extensive research has established a positive correlation between local biodiversity (*e.g.*, species richness and phylogenetic dissimilarity) and multiple functions across various ecosystems and biomes, often regulated by abiotic conditions like climate (Manuel Delgado- Baquerizo et al., 2016; Gonzalez et al., 2020; Jing et al., 2015; Hua Li et al., 2021). For instance, a recent study demonstrated that aridity modulates the relationship between *α*- diversity and soil multifunctionality, with microbial richness showing a stronger positive association with multifunctionality than plant richness in more arid regions (W. G. Hu et al., 2021). This positive relationship is primarily attributed to the selection effect and/or complementary effect among species (Tilman, Isbell, & Cowles, 2014). However, *α*-diversity alone does not sufficiently capture the multiscale nature of biodiversity, which may underestimate the importance of variation of local species assemblages within a heterogeneous environment (*i.e.*, *β*-diversity), especially in sustaining a high multifunctional level concerning the spatiotemporal trade-offs between functions (Sasaki et al., 2022; van der Plas, Hennecke, Chase, van Ruijven, & Barry, 2023).

As a proxy of *β*-diversity, the dissimilarity coefficient (*e.g.*, Jaccard and Sørensen indices) can be partitioned into individual components, such as richness difference and species replacement, which relate to the number of available niches and the influence of ecological gradients on community structure, respectively (Legendre, 2014). The relative contributions of these components to multifunctionality are largely scale-dependent and vary across different ecosystems or biota (Albrecht et al., 2021; Karp et al., 2012; Winfree et al., 2018). In arid environments, species sorting is expected to have a stronger influence on community assembly and the functional effect of *β*-diversity due to their patchy spatial configuration (Guo et al., 2023; Hua Li et al., 2016; van der Plas et al., 2023), highlighting the potential role of species identity and replacement on the variation of multifunctionality (*i.e.*, *β*- multifunctionality) and the maintenance of functions across landscapes. Despite this, the majority of biodiversity-ecosystem function relationship (BEF) studies were emerging from the perspective of single *α*-diversity and its linkage with local functional levels (Manuel Delgado-Baquerizo et al., 2016; W. G. Hu et al., 2021; Soliveres et al., 2016). It would hamper our mechanistic understanding of processes underlying the observed patterns of spatial variation in soil communities and functions along the gradient of climatic transects (A. S. Mori, Isbell, & Seidl, 2018; Rolls et al., 2023).

To address these concerns, we utilized a multitrophic and self-sustainable model community, biological soil crust (biocrust) (Matthew A. Bowker et al., 2014; Garcia-Pichel, 2023), to examine the effects of microbial *α*-diversity on soil multifunctionality within local habitats and those of *β*-diversity components on the spatial variation of multifunctional levels cross a regional gradient of multifaceted precipitation regime (**Supplementary Note 1**).

Biocrusts dominate the topsoil landscape in global drylands (∼ 12% of Earth’s terrestrial surface) with sparse high plants and establish a relatively stable environment against ambient stresses (**Fig. 1a**) (Lange & Belnap, 2016; Weber et al., 2022). The compositions of oxygenic phototrophs (*i.e.*, cyanobacteria), heterotrophic bacteria (hereafter bacteria), and eukaryotic fungi in biocrusts were determined by using the high-throughput amplicon sequencing method. Two *α*-diversity metrics (species richness and mean phylogenetic distance) and the individual *β*-diversity components (richness difference and species replacement) were then quantified. We also estimated the multifunctional importance of each species in consideration of functional redundancy (Hua Li et al., 2021). Moreover, a total of 12 key storage and activity variables were measured to depict the level of soil multifunctionality, reflecting the performance of ecosystems on soil fertility, nutrient cycling, primary production, and climate regulation (Garland et al., 2020).

**Figure 1.**
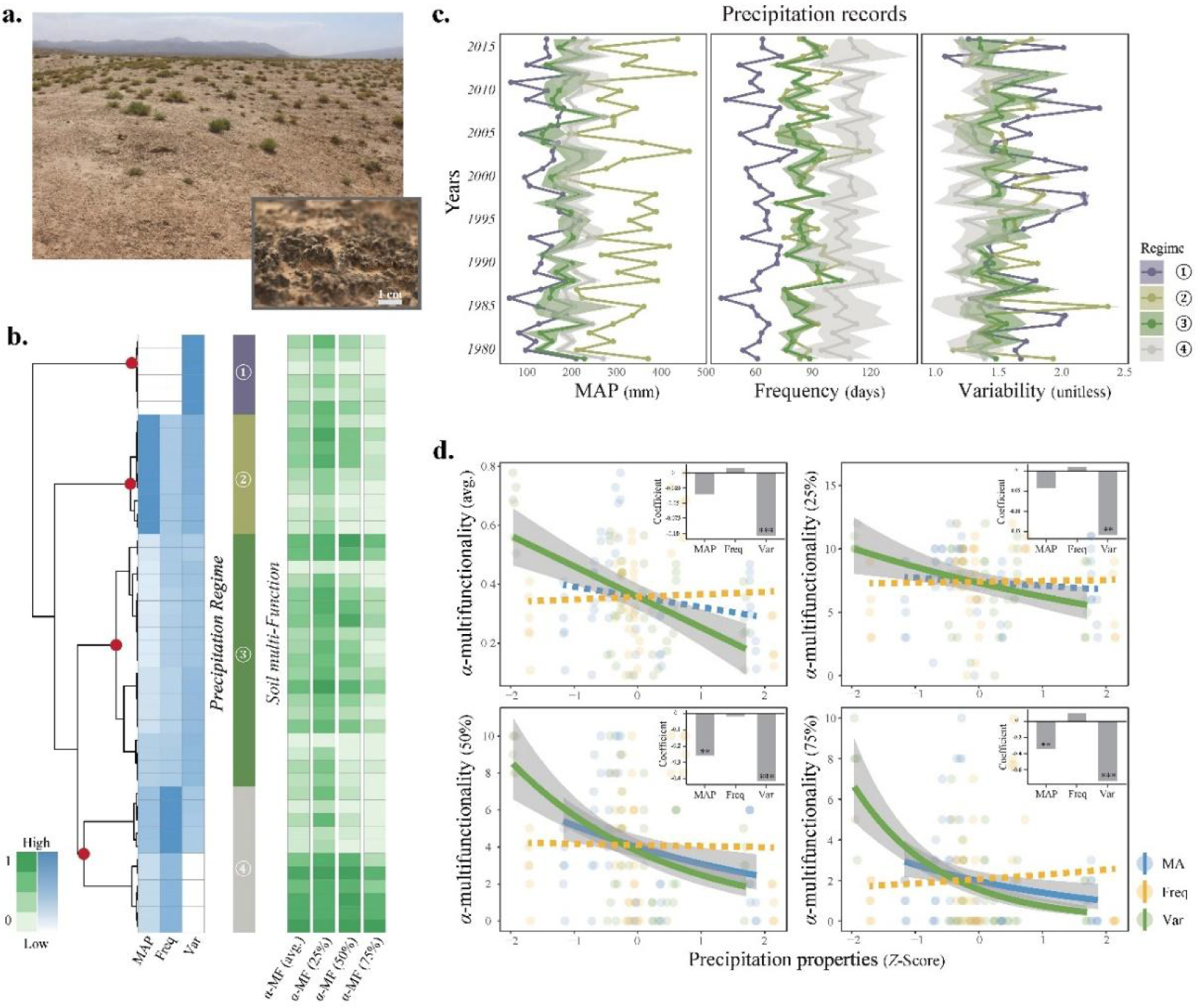
Historical precipitation regime across the study areas in the drylands of northwestern China and its impact on soil *α*-multifunctionality. (**a**) A view of the typical habitat dominated by biocrusts with sparse high plants; (**b**) Clustering tree diagram of study sites based on the dissimilarity of multi-year average precipitation characteristics. The four resulting clusters indicated by dark red points represent regions with distinct precipitation regimes. Corresponding local multifunctionality levels for each site (averaging and threshold- based methods) are displayed on the right panel; (**c**) Annual changes in mean annual precipitation (mm), frequency (days), and variability (unitless) of four regions from 1979 to 2016, shown with respective standard deviation between sites; (**d**) Causal associations between three precipitation characteristics and soil *α*-multifunctionality, fitted by generalized linear models (GLMs). The linkages of averaging [avg., family = gaussion (link = ‘identity’)] and threshold-based *α*-multifunctionality at 25%, 50%, and 75% [family = poisson (link = ‘log’)] are evaluated, respectively. Non-significant relationships (*p* > 0.05) are denoted by dashed lines. The bar chart in the upper right represents the regression coefficients from the GLMs. *α*-MF: local *α*-multifunctionality; MAP: mean annual precipitation amount; Freq: mean annual precipitation frequency; Var: annual daily precipitation variability.

By conducting a cross-regional investigation of biocrusts in northwestern China, an area highly susceptible to perturbations, we aim to unravel the underlying processes of multitrophic responses in soil microbial communities to changes in precipitation, including its amount, frequency, and variability. This will enhance our understanding of the cascading effect of climate change on soil multifunctionality in drylands. We hypothesize that: (**i**) in addition to MAP, various features of the precipitation regime significantly influence multiple functions in biocrusts; (**ii**) given their ecological characteristic relevance, a joint framework incorporating *α*- and *β*-diversity of multitrophic microbes could provide an in-depth elucidation of the regulatory role of biodiversity in the relationship between climate change and ecosystem functioning. Given the representativeness of our research model system and the climatic area studied, we anticipate that the results will underscore the importance of multiple facets of the precipitation regime in global climate prediction modeling and encourage the application of biodiversity theory in ecological restoration practices in drylands worldwide.

## Materials and Methods

### Field sampling and measurement of climate and edaphic variables

In August 2020, we collected biocrust samples from 45 study sites located in the drylands of northwestern China, spanning the Ulanbuh Desert, Hobq Desert, Tengger Desert, and Hexi Corridor (latitude: 37∼41 °N, longitude: 100∼110 °E, see **Supplementary Note 1**). These locations encompass a range of mean annual temperature from 5.4∼9.2 ℃ and MAP from 131∼315 mm. At each site, we randomly collected the upper biocrusts in triplicates from at least three 1×1 m plots. The samples from each site were then thoroughly homogenized and air-dried for subsequent analyses. Further details on the sampling and treatment can be found in our previous work (Guo et al., 2023).

We utilized a high-resolution (30 arc sec, ∼1 km) global downscaled daily precipitation dataset, sourced from the Climatologies at High Resolution for the Earth’s Land Surface Areas (CHELSA: https://chelsa-climate.org/). Using the coordinates of the sampling sites, we extracted daily precipitation data for the period from 1979 to 2016. We then calculated three parameters of multi-year normals to characterize the precipitation regime: mean annual precipitation (*MAP*, mm) to quantify the average precipitation levels, mean annual precipitation frequency (*Freq*, days) to measure the frequency of precipitation events, and coefficient of daily variance to assess the precipitation variability (*Var* = *Standard deviation* / *Mean value*, unitless), based on 0.1-mm and 1.0-mm thresholds for effective daily precipitation, respectively. We also evaluated three key edaphic variables of biocrusts, including the ratio of total organic carbon to total nitrogen (C:N ratio), salinity, and soil pH, which are closely associated with the distribution and assembly of soil microorganisms (Chu et al., 2016; Ni et al., 2021; Rath, Fierer, Murphy, & Rousk, 2019). The measurements of the C:N ratio and soil pH were consistent with the method described in the reference (Guo et al., 2023), and salinity (μmol·g^-1^) was calculated as the sum of the concentrations of related ions (see **Supplementary Note 1**).

### Soil functions and multifunctionality assessment

We evaluated 12 key individual variables related to soil functions in dryland biocrusts: (**i**) chlorophyll-*a* concentration (Chl-*a*, μg·g^-1^), as an indicator of primary productivity; (**ii**) soil fertility variables, including total organic carbon (TOC, mg·g^-1^), total nitrogen (TN, mg·g^-1^), and total phosphorus (TP, mg·g^-1^); (**iii**) available nutrients (NH^4+^, NO ^-^, and PO4^3-^, μg·g^-1^); (**iv**) the activity of enzymes (*β*-glucosidase, sucrase, and alkaline phosphatase), acting as proxies for metabolic ability; and (**v**) water-holding capacity (WHC, w/w) and extracellular polysaccharides content (EPS, μg·g^-1^), which reflect the potential of climatic regulation. The methods for measuring Chl-*a*, TOC, TN, TP, and WHC were specificated previously (H. Li et al., 2020). The contents of NH^4+^, NO ^-^, and PO_4_ ^3-^ were analyzed using ion chromatography (Dionex™ ICS-5000+, Thermo Fisher Scientific, USA). EPS was quantified using the phenol sulfuric acid assay (Dubois, Gilles, Hamilton, Rebers, & Smith, 1956), with glucose as the standard (see **Supplementary Note 1**). Enzyme activity was determined following the extraction and determination procedures provided in the enzyme activity kit (Comin Biotechnology, Suzhou, China).

To take account of potential trade-offs among different soil functions, we calculated Spearman’s correlation coefficients for each pair. In the combinations of 12 functional parameters, about 30 out of 66 pairs exhibited significant positive correlations (|*r*| > 0.6, *p* < 0.001). while one pair showed a negative correlation (**Supplementary** Fig. 1). To standardize the functions with various dimensions, min-max normalization was applied to bring all functional metrics to a unified scale from 0 to 1. Soil *α*-multifunctionality (*i.e.*, local multifunctional level) was then assessed using both averaging and multiple threshold methods (Byrnes et al., 2014; Maestre et al., 2012). In brief, averaging-based *α*-multifunctionality was obtained by calculating the arithmetic mean of the 12 standardized soil functions for each site; and threshold-based *α*-multifunctionality was determined as the count of individual functions exceeding given thresholds of the maximum observed value of each function. We gained the maximum value of each function as the average of the top five sites, to avoid the bias of outliers. BEF relationships were examined at three functional thresholds, 25%, 50%, and 75%, respectively. For *β*-multifunctionality, the multifunctional dissimilarity was measured as the Euclidean distance of the site pair based on the matrix of standardized individual functions (hereafter distance-based *β*-multifunctionality). Meanwhile, for a certain pair of study sites, we also introduced the asynchrony-based *β*-multifunctionality as the number of functions that just exceed a suite of given thresholds (*e.g.*, 25%, 50%, and 75%) of maximum values in one site rather than in the other one, which indicates spatial asynchrony of functionality in a metacommunity (S. Wang, Lamy, Hallett, & Loreau, 2019).

### Estimation of multitrophic local subcommunities in biocrusts

To evaluate the community traits of soil microorganisms in biocrusts, high-throughput amplicon sequencing was employed on an Illumina MiSeq PE300 platform. Bacterial universal primers 338F (5’-ACT CCT ACG GGA GGC AGC AG-3’) and 806R (5’-GGA CTA CHV GGG TWT CTA AT-3’) were used to amplify 16S rRNA gene segments of prokaryotes (H. Mori et al., 2013). For fungi, the ITS region was amplified using primers ITS1F (5’-CTT GGT CAT TTA GAG GAA GTA A-3’) and ITS2 (5’-GCT GCG TTC TTC ATC GAT GC-3’) (Adams, Miletto, Taylor, & Bruns, 2013). As the predominant primary producers in biocrusts, we further estimated the compositions of cyanobacteria by the group- specific primer pair CYA359f (5’-GGG GAA TYT TCC GCA ATG GG-3’) and CYA781r (a: 5’-GAC TAC TGG GGT ATC TAA TCC CAT T-3’; b: 5’-GAC TAC AGG GGT ATC TAA TCC CTT T-3’) to amplify the V3-V4 region of 16S rRNA gene (Nübel, Garcia-Pichel, & Muyzer, 1997).

Following quality inspection and redundancy reduction, amplicon reads were denoised using the unoise3 algorithm to yield zero-radius OTUs (zOTUs, *i.e.*, species) (Edgar, 2018). Taxonomic information was annotated using the RDP algorithm with a threshold set at 0.8. The SILVA v132 database was used for taxonomic annotation of bacteria and cyanobacteria, while the UNITE 8.0 database was used for fungal annotation. According to taxonomic annotation, species assigned to Cyanophyta or chloroplasts were declined from the bacterial dataset amplified by primers 338F/806R, resulting in a subset of heterotrophic soil bacteria (hereafter referred to as bacteria). For the cyanobacterial dataset, eukaryotic chloroplasts and other non-cyanobacterial species were also excluded from subsequent analyses. To minimize disturbance from invalid rare replacement, species with at least 10 reads occurring in more than three sites were selected. The multitrophic datasets were then resampled to control for sampling effort, resulting in 16,463 bacterial, 16,132 cyanobacterial, and 50,921 fungal reads per site. This process yielded 9,526, 1,137, and 3,672 zOTUs for heterotrophic bacteria, oxygenic photoautotrophs (*i.e.*, cyanobacteria), and eukaryotic fungi, respectively.

We measured two metrics of *α*-diversity, namely species richness (zOTU number per site) and mean phylogenetic distance (MPD) among all pairwise species within a given community. In brief, species richness was calculated using the function *diversity* of the *R* package *vegan* v2.5-7. For the MPDs of the abovementioned microbial subcommunities, three maximum- likelihood phylogenetic trees were first constructed with 1,000 bootstraps in IQ-TREE v2.0.3, respectively. The abundance-weighted MPD was then computed by the function *ses.mpd* in the *R* package *picante* v1.8.2, with 999 randomizations and 1,000 iterations of null models (null.model = ‘frequency’) to obtain a standardized effect size (SES) of MPD. General linear models (GLMs) were used to examine the associations between *α*-diversity and local multifunctionality (*α*-MF), both with and without the inclusion of abiotic variables to account for their potential influences. To mitigate multicollinearity among abiotic variables (see **Supplementary** Fig. 2), a principal component analysis (PCA) was priorly performed, and the first five varimax-rotated PCA components (RCs) were extracted using the *R* package *psych* v2.1.9 and included in the multiple regression model, explaining 94.1% of the total variance in the environmental variables (**Supplementary Tab. 1**). When threshold-based *α*- multifunctionality was invoked as the response variable in the models, the estimated values in GLMs (Poisson regression) represented the exponential coefficients of predictors, and the coefficient of determination was calculated as Nagelkerke’s *R*^2^ (Nagelkerke, 1991).

### Partitioning of total β-diversity and evaluation of diversity effects

Spatial *β*-diversity was estimated using the pairwise presence/absence-based Sørensen dissimilarity index (Podani family; *D*_S_), calculated as *D*_S_ = (*b* + *c*) / (2*a* + *b* + *c*), where *a* represents the number of species shared by two sites, and *b* and *c* denote the numbers of species occurring only in one of the paired sites, respectively. *D*_S_ was then decomposed into richness difference [*R*_diff_ = |*b* - *c*| / (2*a* + *b* + *c*)] and species replacement [*Repl* = 2 × min(*b*, *c*) / (2*a* + b + c)] components (using the function *beta.div.comp* in the *R* package *adespatial* v0.3-23), which allows for a more detailed understanding of the mechanisms driving community dissimilarities (Legendre, 2014). Then, we evaluated the effects of climatic, geographic, and edaphic variations on the total *β*-diversity and its components, as well as the effects of both abiotic and *β*-diversity predictors on distance-/asynchrony-based *β*- multifunctionality, by using multiple regression analysis. In this process, geographic distance was declined from the modeling because of its high correlation with the variance of MAP between sites (*i.e.*, ΔMAP) and lower performance by checking the Akaike Information Criterion (AIC) and adjusted *R*^2^ values of sub-models (**Supplementary** Fig. 3 and **Supplementary Tab. 2**). To gain a measure of uncertainty for the evaluation of the diversity effect, a bootstrap procedure was implemented by resampling a subset of 60% of all sites without replacement (*n* = 1,000 replicates) using the *R* package *boot* v1.3-28. The mean value, as well as the 50% and 95% confidence intervals, for each predictor were then calculated.

The piecewise structural equation model (pSEM) was employed to further assess the cascading associations between microbial *β*-diversity components (*Repl* and *R*_diff_) and *β*- multifunctionality, while simultaneously accounting for the variances of key environmental factors such as geography (ΔElevation), precipitation regime (ΔMAP, ΔFreq, and ΔVar), and soil properties (ΔpH, ΔSalinity, and ΔC:N). An *a priori* schematic diagram of pSEM was constructed before analysis, with the mechanism of each putative path validated by previous references (**Supplementary** Fig. 4). The Fisher’s *C*-test (0.05 < *p* < 1.00) along with the AIC and Bayesian Information Criterion (BIC) were used to compare nested models and confirm the overall goodness of fit. Ultimately, those significant (*p* < 0.05) and influential (|λ| > 0.2) paths were included for visualization. The pSEM analysis was conducted in the *R* package *piecewiseSEM* v2.1.2.

### Multifunctional importance and environmental preference

The multifunctional importance of microbial taxa at the species level was evaluated using randomization tests in the Fortran 95 program *Impact* (Gotelli, Ulrich, & Maestre, 2011). Both presence-absence and relative abundance datasets were used to calculate species-specific values of multifunctional importance, considering the averaging-based *α*-multifunctionality.

For the presence-absence matrix, the difference in multifunctionality was measured when the target species was present or absent in a certain site pair. For the abundance matrix, the rank correlation coefficient between each species’ relative abundance and soil multifunctionality was calculated. We conducted randomization (999 permutations) to reassign observed values across different sites and repeat the null modeling for each species, obtaining the SES value of multifunctional importance (Hua Li et al., 2021). It indicated the degree to which observed values deviate from the null distribution generated by random simulation. A species with SES > 1.96 or < -1.96 was supposed to have a significant positive or negative effect on local multifunctionality in biocrusts, respectively, while |SES| < 1.96 of species meant a non- significant importance. According to the category of species functionality, we also computed the relative contribution of species to the total *β*-diversity of pairwise sites by using the similarity percentage (SIMPER) analysis (Clarke, 1993), thereby determining the cumulative contributions of different categories to the total variance of the metacommunity (**Supplementary** Fig. 5).

To examine whether the distribution of species with varying multifunctional importance follows a pattern along abiotic gradients, we calculated the extent of environmental preference for each species by averaging each abiotic variable weighted by the local abundance across all sites (Jiao et al., 2022). We then fitted piecewise linear regressions between species’ multifunctional importance and their environmental preferences using the *R* package *segmented* v1.6-0. Threshold values were evaluated to identify breakpoints where the slope exhibited significant changes. Model selection was performed based on AIC and adjusted *R*^2^ values, which suggested two breakpoints for optimal modeling performance.

## Results

### Precipitation regime and the impact on soil multifunctionality

An analysis of a daily precipitation dataset spanning 38 years revealed the historical precipitation patterns in the arid and semi-arid regions of northwestern China. Four distinct clusters emerged based on three parameters of multi-year average precipitation records (MAP, Freq, and Var), each representing a unique precipitation regime. These parameters were not entirely covariant, suggesting their independence and emphasizing the need to consider multifaceted precipitation characteristics (**Fig. 1b**).

Overall, annual precipitation demonstrated fluctuating dynamics, with precipitation variability likely to exhibit a greater degree of oscillation compared to MAP and frequency (**Fig. 1c**). In recent decades, regions with differential regimes of precipitation (Regimes 1 to 4) largely experienced heterogeneous historical precipitation changes. For instance, the sites within Regime 2 underwent the highest fluctuation of MAP with a relatively moderate change in frequency, while sites in Regime 3 had lower fluctuations in all precipitation facets than others (**Supplementary Note 1**). These findings were corroborated by analyses using a 1-mm threshold for effective precipitation, aligning with results obtained using a 0.1-mm threshold (**Supplementary** Fig. 6).

Regardless of the method used (averaging or multiple thresholds), a significant negative correlation was observed between precipitation variability and soil local *α*-multifunctionality (**Fig. 1d**). Nonetheless, MAP showed a significant negative correlation with threshold-based *α*-multifunctionality when moderate and high levels of functioning were desired (50∼75%). For precipitation frequency, no considerable association was detected with *α*-multifunctionality. In the context of *β*-multifunctionality, a similar result was observed where the difference in precipitation variability (ΔVar) significantly explained a larger portion of the variance in soil multifunctionality between sites (*p* < 0.001), compared to MAP and precipitation frequency (**Supplementary Tab. 3**).

### α-Diversity effects on local multifunctionality

At the local scale, soil pH and salinity served as the most significant predictors for a variety of multitrophic *α*-diversity indices (richness and ses.MPD) in biocrusts (**Supplementary** Fig. 7). We found that fungal *α*-diversity was more closely related to abiotic factors than that of heterotrophic bacteria and oxygenic phototrophs (cyanobacteria), with further influences from geographic properties (longitude and latitude) and precipitation variability. In contrast, bacterial MPD seemed to be unaffected by any measured abiotic variables, indicating a relatively stochastic pattern of phylogenetic signal.

Significant correlations were broadly observed between species *α*-diversity and local soil multifunctionality (using both averaging and threshold-based methods) across multitrophic microbiota in biocrusts (**Fig. 2a, b**). However, after adjusting for co-varying abiotic factors, the richness of heterotrophic bacteria and fungi, along with cyanobacterial MPD, retained their positive effects on soil multifunctionality. This suggests a differentiated underlying mechanism across various trophic levels in biocrusts that enhances soil functioning (**Supplementary Tab. 4**). Furthermore, our analysis revealed that regardless of the method used (averaging or threshold-based) or the extent of achieved multifunctional level considered, cyanobacterial MPD (estimated coefficient β: 0.024∼0.382) and fungal richness (β: 0.029∼0.347) emerged as the most influential drivers of microbial *α*-diversity components on local multifunctionality of biocrusts, even after controlling for abiotic confounding factors (**Fig. 2c** and **Supplementary Tab. 1**).

**Figure 2.**
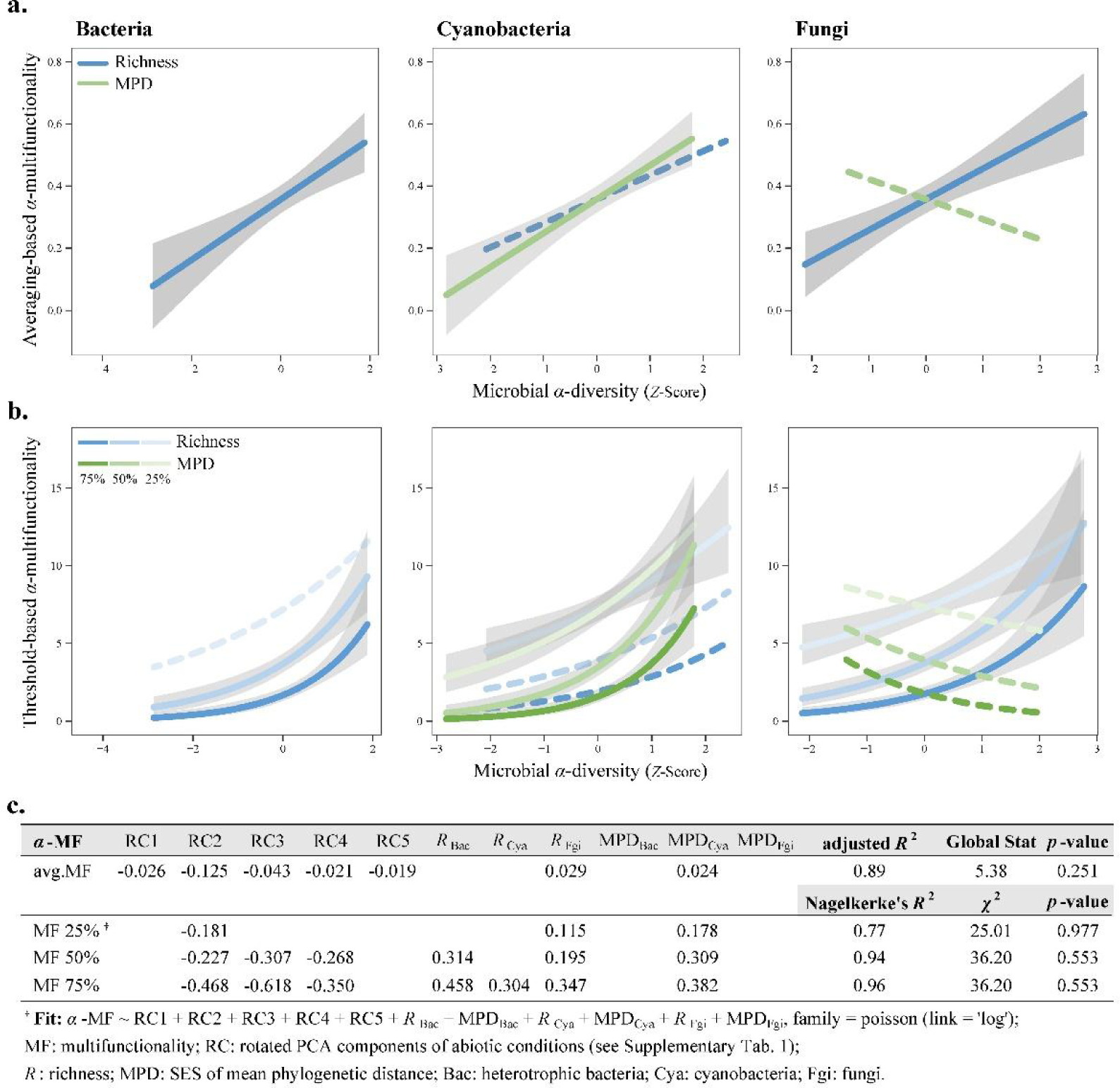
Diversity effects of multitrophic microbiota in biocrusts on soil *α*-multifunctionality. The relationships between each *α*-diversity index [richness and standardized effect size of mean phylogenetic distance (ses.MPD)] and local multifunctionality are tested using averaging (**a**) and multiple threshold approaches (**b**) by fitting generalized linear models (GLMs), respectively. Three thresholds from low to high (25%, 50%, and 75%) are chosen and represented by a color gradient. Solid lines indicate significant relationships even when considering abiotic confounding factors, while dashed lines represent the associations that turn into non-significant in multiple regressions. (**c**) Along with pairwise GLMs, multiple regressions are performed to examine the potential influences of abiotic conditions and multitrophic interaction on diversity effects. Complete models are built for both averaging and threshold-based *α*-multifunctionality, including richness and ses.MPD of three subcommunities and five abiotic rotated principal components as predictive variables. A stepwise procedure is employed to obtain the optimal model by checking AICs. The global validation of linear model assumptions is estimated using the *R* package *gvlma* (Global Stat), and a Chi-Square (*χ*^2^) goodness of fit test is used for Poisson regression.

### Spatial richness difference and species replacement of microbiota

At the cross-regional scale of drylands, community dissimilarity (*i.e.*, *β*-diversity) of multitrophic levels was primarily driven by species replacement (**Fig. 3**). Specifically, it accounted for 81.96%, 67.67%, and 78.60% of the variance in the total *β*-diversity for bacteria, cyanobacteria, and fungi, respectively. Fungi exhibited significantly higher mean total *β*- diversity and species replacement than the other two subcommunities, with cyanobacteria showing the lowest values (**Fig. 3a, b**, Wilcoxon test *p* < 0.001). Indeed, the relationships between *β*-diversity and geographic distance indicated the highest spatial turnover rates in soil fungi but the lowest rates in cyanobacteria (**Supplementary** Fig. 8).

**Figure 3.**
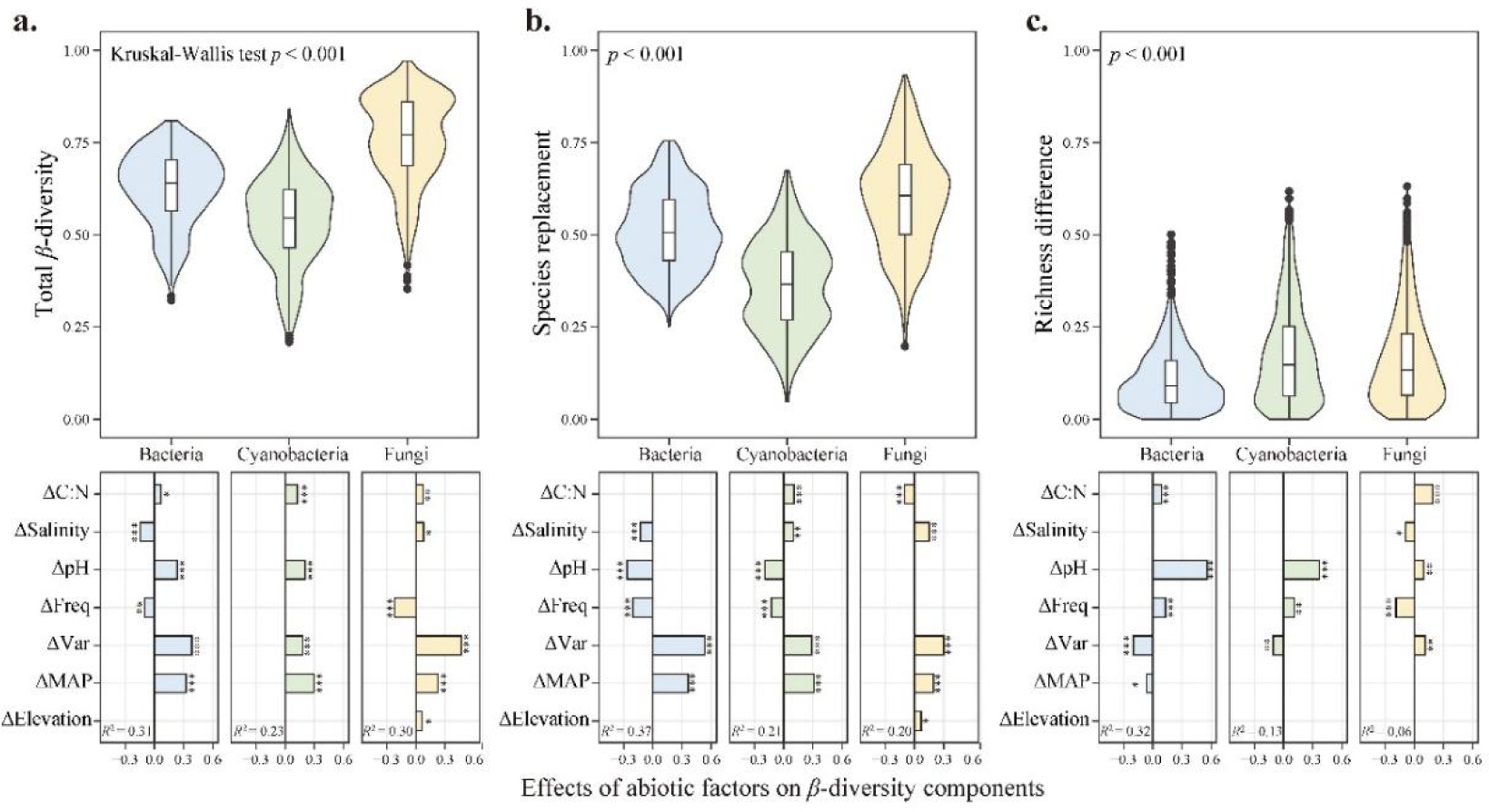
Impact of abiotic heterogeneity on total *β*-diversity and its components. (**a**) Total *β*- diversity (Sørensen dissimilarity) between sites is decomposed into richness difference and species replacement. The significance of inter-group differences among three trophic levels of soil microorganisms is evaluated using the Kruskal-Wallis tests. (**b**) Multiple regression is performed for total *β*-diversity, richness difference, and species replacement, respectively, to explore their responses to changes in abiotic conditions. The symbol Δ represents the spatial variance of a specific abiotic variable between sites, and significance levels are indicated as *** *p* < 0.001, ** *p* < 0.01, and * *p* < 0.05. MAP: mean annual precipitation amount; Freq: annual precipitation frequency; Var: daily precipitation variability.

To further investigate the environmental response of *β*-diversity, we examined the associations between *β*-diversity components and abiotic heterogeneity of site pairs using multiple regression analyses. In general, climatic, edaphic, and geographic dissimilarities explained a sum of 23.27∼31.25% variance in total *β*-diversity, 5.99∼31.84% in richness differences, and 19.76 ∼ 36.72% in species replacements of three trophic subcommunities. We found that the variances in MAP, precipitation variability, and soil pH were the most predominant factors determining spatial community dissimilarities of soil microorganisms in drylands. Nonetheless, as additive components of *β*-diversity, richness difference and species replacement exhibited an underlying bias in their responses to various abiotic dissimilarities. That is, species replacement in all subcommunities was highly susceptible to changes in MAP and variability (ΔMAP and ΔVar), while richness difference was primarily influenced by changes in edaphic conditions, particularly soil ΔpH (**Fig. 3b, c**). Notably, compared to heterotrophic bacteria or fungi, terrestrial cyanobacteria in biocrusts appeared to be less sensitive to changes in the precipitation regime. Moreover, we only included ΔMAP in the models instead of the geographic distance between sites (Geodist), due to their high correlation (*i.e.*, multicollinearity problem). However, when Geodist was employed instead, most models gained higher adjusted *R*^2^ values (**Supplementary Tab. 5**). This suggests that in addition to ΔMAP, other unmeasured variables with spatial autocorrelation or dispersal limitations also contribute to the *β*-diversity patterns of multitrophic subcommunities in the drylands of northwestern China.

### Joint effects of abiotic heterogeneity and β-diversity on multifunctionality

We observed a strong positive correlation between microbial *β*-diversity components and distance-based *β*-multifunctionality, taking into account the combined effects with abiotic heterogeneity (**Fig. 4**). This correlation was predominantly driven by species replacement and displayed consistent patterns across all three subcommunities (standardized coefficient β_Bac_ = 0.51, β_Cya_ = 0.51, and β_Fgi_ = 0.55, **Supplementary Tab. 6**). Meanwhile, changes in soil salinity (ΔSalinity) and precipitation variability (ΔVar) were the most influential factors among the edaphic and climatic variables (β_ΔSalinity_: 0.30∼0.40 and β_ΔVar_: 0.24∼0.34), albeit less than the diversity effects. Moreover, the variance of multifunctionality was also calculated by evaluating the spatial asynchrony across different individual functions. As the required level of multifunctionality increased from 25% to 75%, a significant shift in the relative importance of abiotic versus diversity effects occurred, primarily for heterotrophic bacteria and cyanobacteria (**Supplementary Tab. 7**). We found that precipitation heterogeneity, especially variance in variability (ΔVar), became more considerable in predicting *β*-multifunctionality at higher thresholds (the coefficient β_ΔVar_ increased).

**Figure 4.**
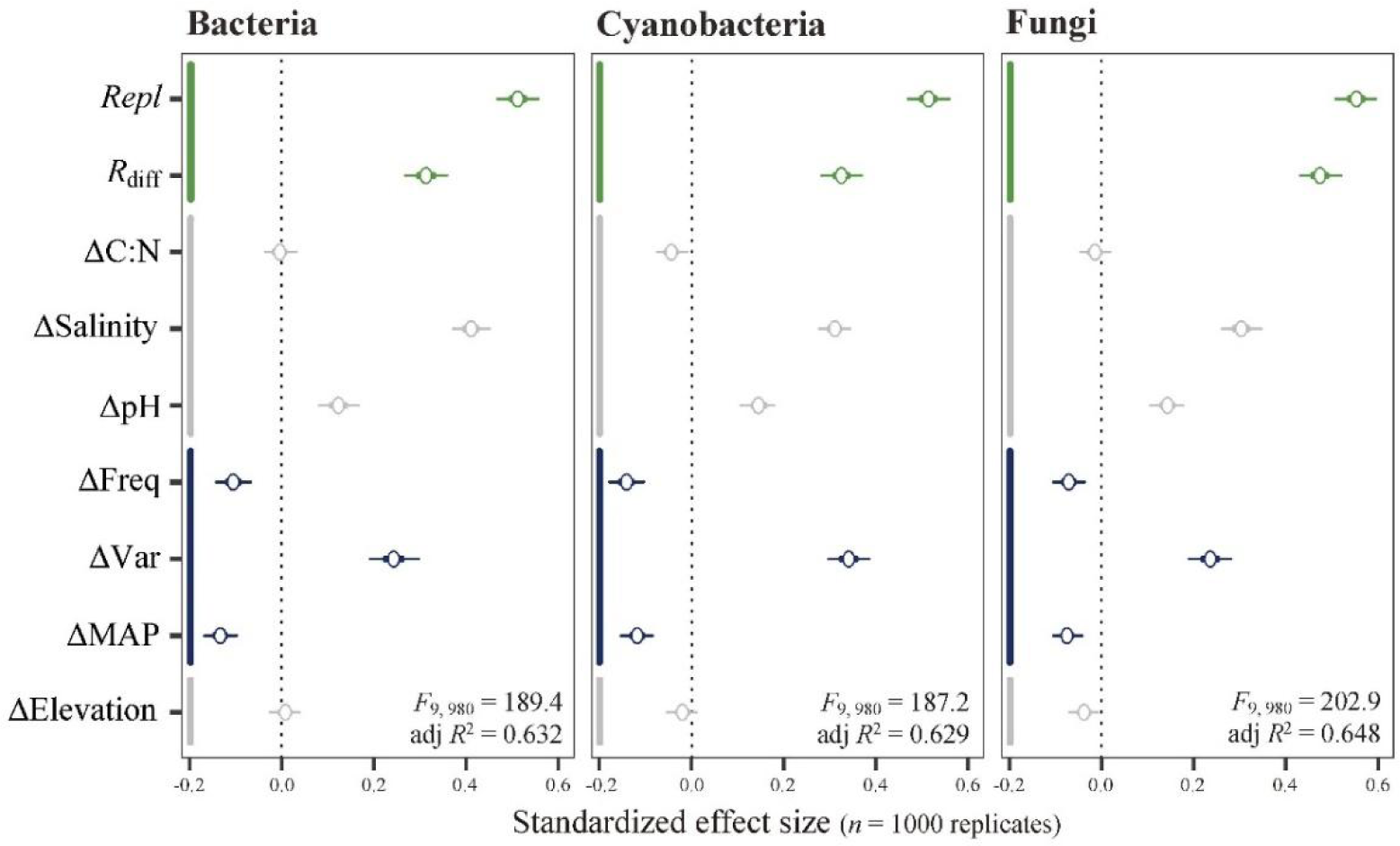
Combined effects of *β*-diversity components and abiotic heterogeneity on variations in soil multifunctionality across different microbial subcommunities in biocrusts. *β*- multifunctionality is quantified as the Euclidean distance of standardized individual functions between sites. Estimated effects of richness difference and species replacement (green), as well as edaphic and geographic variables (gray) and variances in precipitation regime (navy blue), are evaluated by multiple regression. A bootstrap procedure is used to reflect the extent of fluctuation in each estimated effect size of predictors among sites (*n* = 1,000 replications). The mean values (circles), as well as 50% and 95% confidence intervals (thin and thick bars, respectively), are denoted. MAP: mean annual precipitation; Freq: annual precipitation frequency; Var: daily precipitation variability; *R*_diff_: richness difference; *Repl*: species replacement.

Nevertheless, for soil fungi, the richness difference and species replacement maintained their influences at any given level of multiple functions and achieved a balance with abiotic effects at the highest level (β*_R_*_diff_ = 0.25, β*_Repl_* = 0.34, β_ΔVar_ = 0.28, and β_ΔSalinity_ = 0.17).

The causality network between geography, climate, soil environment, microbial *β*- diversity, and multifunctional variance, constructed by pSEM, highlighted a cascading pathway from the precipitation regime to soil multifunctionality, mediated by *β*-diversity, in the drylands of northwestern China (**Fig. 5**). Compared to the direct effect of precipitation heterogeneity, its indirect influences on *β*-multifunctionality (distance-based) through richness difference, species replacement, and variance in soil properties were significantly more dominant across multitrophic microbiota of biocrusts. These indirect influences mainly arose from a cascade of variance in variability (ΔVar) to species replacement (effect size θ_ΔVar *-> Repl*_: 0.29∼0.53) and ΔSalinity (θ_ΔVar -> ΔSalinity_ = 0.54) and ultimately converged on *β*- multifunctionality (θ*_Repl_*: 0.51∼0.55, and θ_ΔSalinity_: 0.31∼0.41, see **Fig. 5**). Furthermore, when we assessed causality using asynchrony-based *β*-multifunctionality with a gradient of thresholds, a shifting weightiness of *β*-diversity, soil properties, and precipitation heterogeneity was confirmed in line with the results of multiple regression (**Supplementary** Figs. 9**∼11** and **Supplementary Tab. 7**). At the low and intermediate levels of functional threshold (25% and 50%), climatic and environmental factors only impacted *β*- multifunctionality via the direct influence mediated by *β*-diversity components of soil microorganisms (θ*_R_*_diff_: 0.32∼0.54 and θ*_Repl_*: 0.38∼0.68, see **Supplementary** Figs. 9**∼10**), reflecting an amplifying effect of *β*-diversity. However, when a higher level of functional threshold was considered, the abiotic predictors ΔVar, followed by ΔSalinity, emerged as having direct functional influences and eventually explained most of the variance in *β*- multifunctionality (*e.g.*, θ*_R_*_diff_ and θ*_Repl_* < 0.20 in heterotrophic bacteria and cyanobacteria at the 75% threshold, see **Supplementary** Fig. 11).

**Figure 5.**
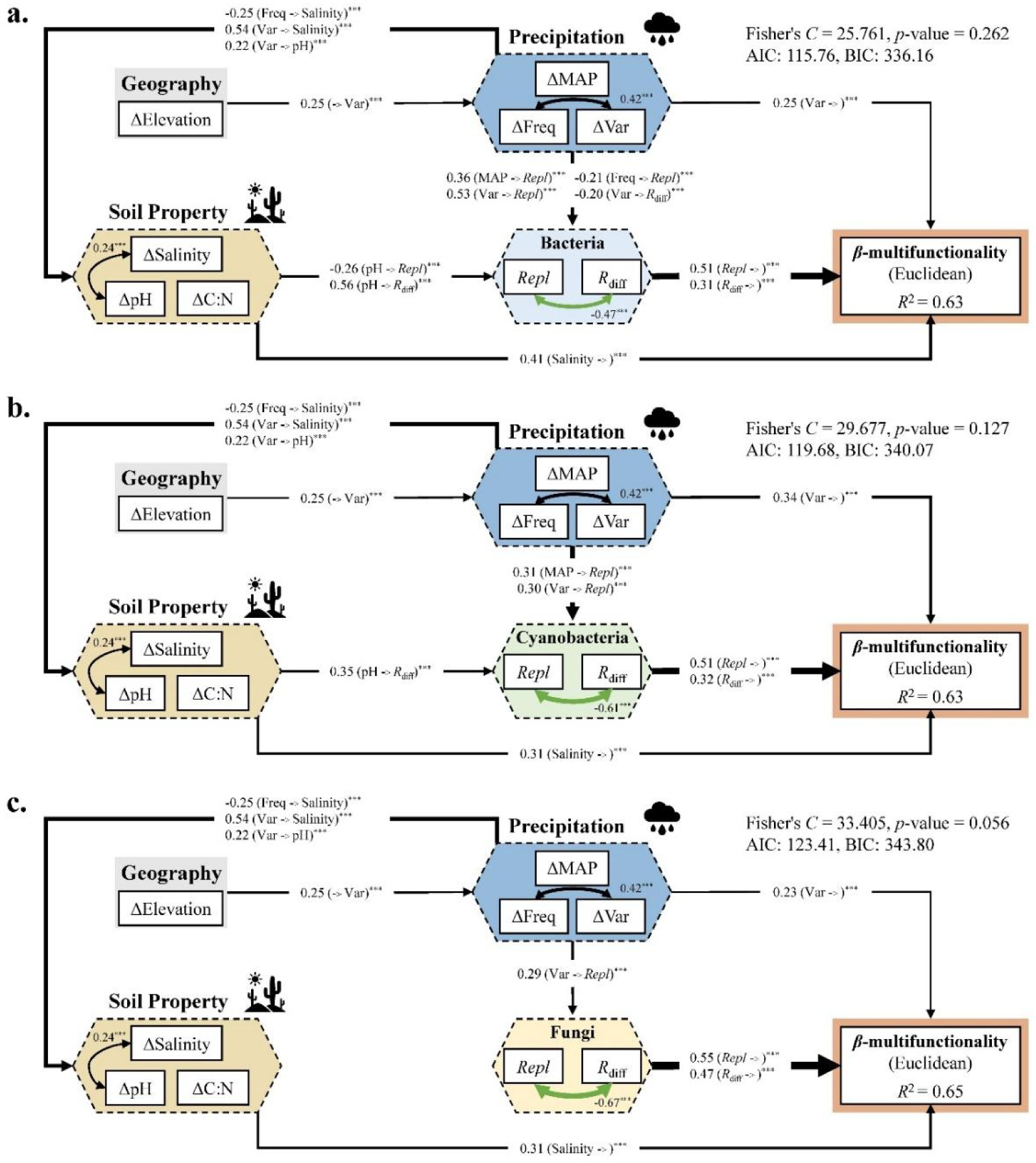
Cascading topology of piecewise structural equation models illustrating the impact of abiotic heterogeneity and multitrophic *β*-diversity components on soil multifunctionality. The connection networks depict the direct and indirect contributions of geographic (Δ Elevation), climatic (ΔMAP, ΔFreq, and ΔVar), and edaphic heterogeneities (ΔpH, ΔSalinity, and ΔC:N), mediated by richness difference and species replacement of heterotrophic bacteria (**a**), cyanobacteria (**b**), and fungi (**c**), to variations in soil multifunctionality. Black arrows indicate significant positive paths and green arrows indicate significant negative paths. The significance of each path is indicated as *** *p* < 0.001, ** *p* < 0.01, * *p* < 0.05, and the number adjacent to the arrows represents the standardized path coefficient. Arrows with an integrated net effect size of each predictor category |λ| ≥ 0.2 are presented, and the width of the arrows is proportional to the net effect size. Fisher’s *C* statistic, *p*-values, Akaike and Bayesian Information Criteria (AIC and BIC) of models are shown as the global test of goodness of fit. MAP: mean annual precipitation; Freq: annual precipitation frequency; Var: daily precipitation variability; *R*_diff_: richness difference; *Repl*: species replacement.

### Microbial climate preferences predict functional importance

At the species level, the SES of multifunctional importance index based on the presence- absence matrix showed that 27.59∼35.97% of soil microorganisms significantly contributed more to the local multifunctionality of biocrusts than expected by chance, therein 21.57∼25.07% of multitrophic microbiota were associated with the enhancement of multifunctionality when they were present (**Supplementary** Fig. 12). Similar results were obtained by using the abundance matrix of species. Among the trophic levels, oxygenic photoautotrophs (cyanobacteria) consisted of a lower proportion of species (64.03%) with non-significant importance than that of heterotrophic bacteria (67.43%) and fungi (72.41%), which to some extent implied a differentiation of functional redundancy in different subcommunities of biocrusts (Wilcoxon rank sum tests, *p* < 0.05).

To understand how the multifunctional significance of species changes with their preferences for various abiotic conditions, we calculated the environmental preference value for each species and established a piecewise linkage with the functional importance index. For precipitation regimes, the species distribution of multifunctional importance exhibited a peak within a range of 180∼200 mm of MAP and 80∼100 days of mean annual precipitation frequency, while rapidly declining at either low or high end of gradients (**Fig. 6a, b**), which can be often analogized by the intermediate distribution hypothesis. However, irrespective of trophic levels, there was a successive reduction in multifunctional importance, particularly at the mid-gradient of precipitation variability around 1.45 (unitless, **Fig. 6c**), indicating a potential accumulation of such low-functioning microorganisms in response to elevated variability. Additionally, we investigated whether the edaphic preferences of soil microbiota could explain their variations in multifunctional importance. The findings demonstrated that species of positive importance were typically prevalent in environments characterized by relatively lower soil pH (∼8.4, unitless) and appropriate salinity (∼7.50 μmol·g^-1^) in drylands (**Supplementary** Fig. 13).

**Figure 6.**
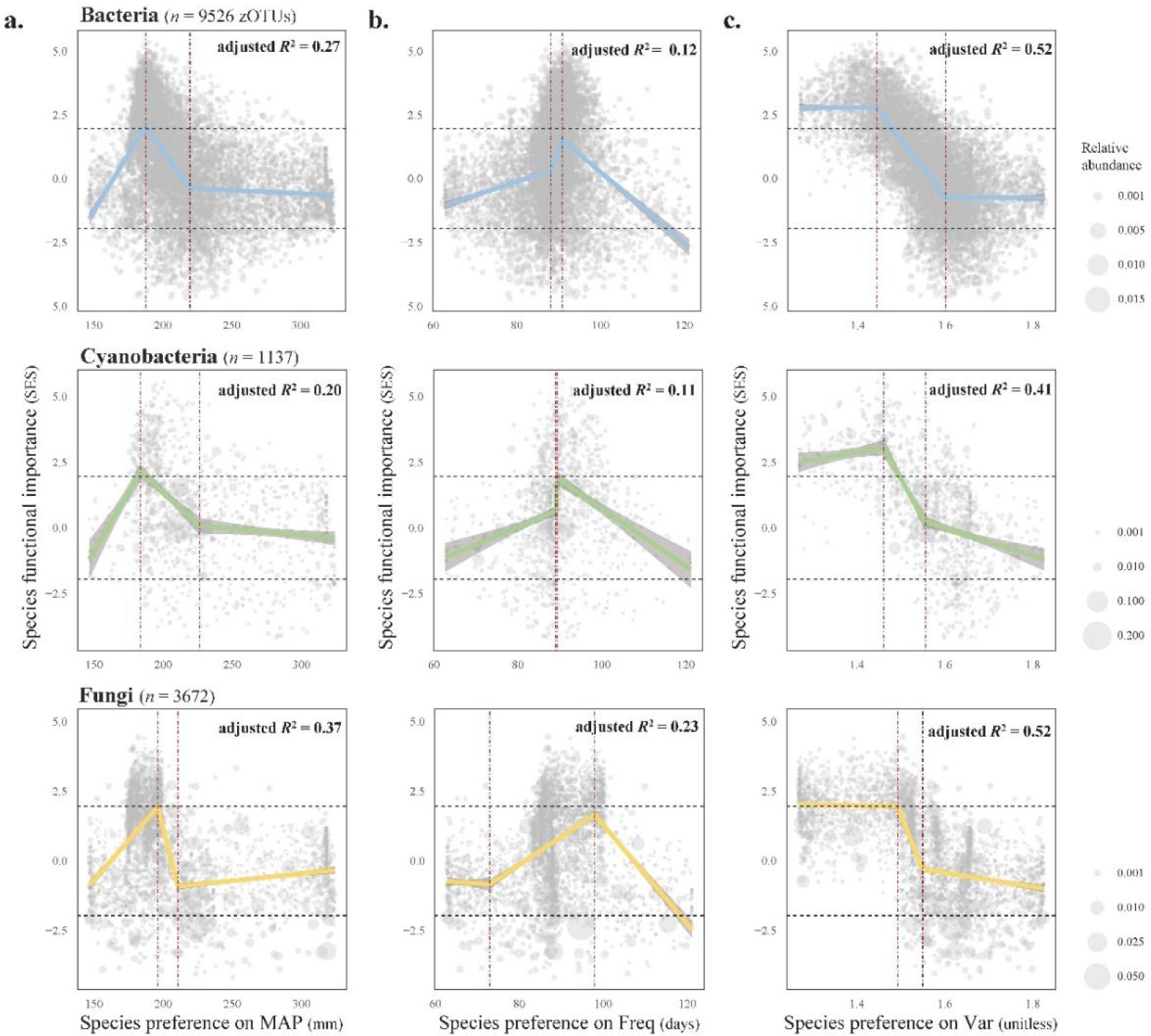
Species preference for precipitation regime predicts their multifunctional importance. Nonlinear responses of multifunctional significance to species preference for mean annual precipitation (**a**), annual precipitation frequency (**b**), and daily precipitation variability (**c**) are described using piecewise regression. Dark red dashed lines indicate the breakpoints of thresholds where the slope shows significant changes. Black dashed lines denote the critical thresholds of the 95% confidence interval (CI) of a null model, where the standardized *Z*-value of multifunctional importance is significantly lower or higher than expected by chance. The grey area of regression lines represents the 95% CI of piecewise regression. Species multifunctional importance is calculated based on the presence-absence matrix. MAP: mean annual precipitation; Freq: annual precipitation frequency; Var: daily precipitation variability.

## Discussion

In this study, we explored the diversity effects of multitrophic microbiota on soil functionality by considering multiple functions at local and cross-regional scales in the drylands of northwestern China. In recent decades, it has been increasingly accepted that high local species richness is crucial for sustaining the simultaneous performance of various ecosystem functions and services (Albrecht et al., 2021; M.A. Bowker, Maestre, & Mau, 2013; Hooper et al., 2012; Wall, 1999). This positive BEF relationship is mainly attributed to the extent of functional dissimilarity between co-existing species (*i.e.*, functional complementarity) or the selection effect of the local species pool (*i.e.*, the dominance of particular highly functioning species) (Loreau & Hector, 2001; Tilman et al., 2014). Nonetheless, emerging studies shed light on the intricate nature of linkages between richness and functionality in different contexts and biomes (M. A. Bradford et al., 2014; Jing et al., 2015).

More recent perspectives propose that the temporarily realized richness of species may not guarantee a positive association with the current level of ecosystem function (Hagan, Vanschoenwinkel, & Gamfeldt, 2021). This is due to a balance between complementarity and selection effects that fluctuate during the process of community assembly. Indeed, after controlling for abiotic covariates, we observed differentiated *α*-diversity-function relationships across different trophic levels of soil microbiota in biocrusts (**Fig. 2**).

Specifically, the local richness of heterotrophs (bacteria and fungi) maintains a significant positive association with soil multifunctionality, whilst oxygenic photoautotrophs (cyanobacteria) exert the positive functional effect through phylogenetic dissimilarity (*i.e.*, MPD) rather than species richness. It is noteworthy that there is an intrinsic difference in the phylogenetic and physiological breadth of members in the three subcommunities, which fulfill relatively broad or narrow spectrums of functional processes. As a result, an elevated functional redundancy [also proposed as *functional similarity* in a multifunctional context, see (Eisenhauer, Hines, Maestre, & Rillig, 2023)] among specialized cyanobacterial species compared to that of the entire heterotrophic bacteria or fungi is predictable and has been verified in previous research (M. Delgado-Baquerizo et al., 2016; A. S. Mori et al., 2016). By contrast, as a proxy for phylogeny-related trait dissimilarity of species, standardized MPD measures the ability of functional resilience within cyanobacterial assemblages against ambient disturbances, exhibiting a stronger explanatory power for the variance in soil multifunctionality (Hua Li et al., 2021).

Multiscale dependence is an inherent characteristic of biodiversity. In this context, *β*- diversity bridges local (*α*-) and larger-scale (*γ*-) biodiversity and sustains multiple functions by species sorting in heterogenous landscapes (A. S. Mori et al., 2018; van der Plas et al., 2023). To enhance our mechanistic understanding of the functional roles of *β*-diversity, we partition the total diversity into richness difference and species replacement components for three multitrophic subcommunities, respectively. In most modeling scenarios, we discover a consistently stronger positive effect of species replacement on the variations in soil multifunctionality than that of richness difference, despite both of them having more significant functional signals than abiotic predictors (**Fig. 4** and **Fig. 5**). This finding has three major ecological implications. First, despite the typically skewed distribution of microbial abundances in soils dominated by a few species (Pedros-Alio, 2012), it is supposed that there are no predominant high-functioning species that outcompete others in the meta-community of our regions. This notably contrasts with the widespread view of terrestrial cyanobacteria that a limited set of filamentous species (*e.g.*, *Microcoleus*, *Leptolyngbya*, and *Scytonema* spp.) undertakes the main ecological roles in biocrust-forming and soil functioning (Belnap, 2002; C. X. Hu & Liu, 2003). However, it is worth noting that we essentially evaluate microbial compositions of biocrusts at a sub-species level (zOTUs at the 100% identity threshold). It certainly highlights the significance of both inter- and intraspecific identity in maintaining the *β*-diversity effect through species replacement (Isbell et al., 2018; Winfree et al., 2018).

Second, as a model system for BEF relationship research (Matthew A. Bowker, Maestre, & Escolar, 2010), biocrusts create a variety of patchy landscapes where heterogeneous spatial niches harbor different subsets of the regional species pool, maximizing their functioning in different patches (*i.e.*, species sorting) (Leibold, Chase, & Ernest, 2017). Consequently, abiotic heterogeneity of biocrusts ensures the strong effects of both richness difference and species replacement on the variance of soil multifunctionality in drylands. The relative importance of two *β*-diversity components may shift in response to changes in environmental heterogeneity or depending on the ecosystem types considered (Albrecht et al., 2021). Third, the diversity-mediated biotic effect of soil multitrophic microbiota is likely to amplify the ecological consequences of climate change (see **Fig. 5**). Whether considering dissimilarity or spatial asynchrony of multifunctionality, the direct effect of abiotic heterogeneities is less influential than the indirect effect via *β*-diversity components. This pattern becomes particularly evident when we evaluate the functional asynchrony at low and intermediate thresholds (25∼50%), wherein abiotic factors merely exert indirect effects on the variance of multifunctionality, relying on microbial diversity (**Supplementary** Figs. 9**∼10**). However, if high levels of multiple functions are desired at heterogeneous landscapes, *β*-diversity- mediated influences may collapse and the heterogeneity of environmental conditions tends to set the upper boundary of functional potential (**Supplementary** Fig. 11) (van der Plas et al., 2016).

Our analyses show that the spatial heterogeneity of the precipitation regime largely drives species replacement in all three subcommunities, meanwhile, richness difference, especially in heterotrophic bacteria, primarily arises due to variations in soil pH (**Fig. 3**). This is aligned with a previous study by Evans & Wallenstein (2014), who found that soil microbes can alter their community composition (enhancing temporal *β*-diversity) and change the distribution of moisture-related ecological strategies (based on species identity) when exposed to intensified rainfall patterns (S. E. Evans & Wallenstein, 2014). In contrast to fungi, soil pH demonstrated its ability to exclusively structure bacterial communities and influence their richness in liming experiments (Rousk et al., 2010; Seaton et al., 2020). Therefore, we speculate that edaphic conditions, such as soil pH, play a crucial role in controlling local species coexistence by impacting niche availability and partitioning in biocrusts, while large-scale climatic factors like precipitation regime seem to be more prominent in driving microbial turnover along gradients (Gonzalez et al., 2020; Stein, Gerstner, & Kreft, 2014). In addition, we observed a faster spatial turnover rate for fungi (**Supplementary** Fig. 8), likely due to their rapid adaption of community composition to environmental changes (Voriskova, Elberling, & Prieme, 2019). In contrast, cyanobacterial lower turnover rate implies greater resistance to precipitation fluctuations, possibly due to their survival strategy that anticipates desiccation (Rajeev et al., 2013; Xu et al., 2021). However, although heterotrophic bacteria and fungi exhibit higher turnover rates, they are largely comprised of exchanges between functionally nonsignificant species (see **Supplementary** Fig. 5). This could explain why cyanobacteria, despite their lower turnover, effectively support the variance in multifunctionality across regions. More importantly, as expected, our results indicate that the variability, rather than the amount or frequency of multi-year average precipitation, seems to be the most significant abiotic driver of the variance in soil multifunctionality, which may impact nutrient enrichment and soil aggregate stability (Yao et al., 2020). Furthermore, the functionality of multitrophic microbiota in biocrusts, as assessed by their multifunctional importance index, shows a uniform sensitivity against the gradient of precipitation variability on a large scale, thus high variability can directly lead to a convergence of species with low functional importance (**Fig. 6**), thereby altering the functional composition of communities. Given its precedence over changes in other traditional precipitation properties (Ren, Sha, Shi, & Liu, 2021), precipitation variability caused by climate change could subtly impair the ability of dryland ecosystems to maintain multiple functions before it becomes noticeably apparent.

## Conclusion

Human beings are currently confronting the widespread impacts of climate change on the delivery of ecosystem functions and services, a main characteristic of the Anthropocene era. Although extensive attention condenses on the alteration in aridity and its direct influences on biodiversity and ecosystem functionality, the multifaceted properties of precipitation regimes and their indirect effects remain largely unknown. To address this, we used a BEF framework to examine the diversity effects of multitrophic microbiota and abiotic influences on soil functionality across local and cross-regional scales in the drylands of northwestern China. Our findings highlight the importance of both *α*- and *β*-diversity in maintaining soil multifunctionality. Nevertheless, the underlying mechanisms are differentiated. The local richness of heterotrophs but the phylogenetic dissimilarity of photoautotrophs exerted positive functional effects on soil multifunctionality. We also found that *β*-diversity, particularly replacement based on species identity, played a significant role in supporting changes in soil multifunctionality. It is noteworthy that precipitation variability, rather than MAP, emerged as the predominant driver of multifunctionality, potentially leading to a convergence of less beneficial species in soil communities. Most importantly, the results underscore the cascading impacts of precipitation variability on dryland ecosystems via the amplifying effect of microbial diversity, which may induce a ‘silent’ breakdown of those vulnerable arid and semiarid environments. Despite the absence of tested experimental causality, our study still contributes to an in-depth understanding of the underlying mechanisms of how climate change impacts dryland multifunctionality in the real world and ultimately promotes the management of ecosystem restoration.

## Acknowledgments

The research is supported by the National Natural Science Foundation of China (Grant Nos. 41877419 and 32370125) and the Natural Science Foundation for Distinguished Young Scholars of Hubei Province (No. 2022CFA105). We are grateful for sampling assistance from Da Huo and technical support from the Analysis and Testing Center of IHB, and the sequencing and statistical analyses are assisted by the Supercomputing Center of CAS, Wuhan Branch.

## Appendixes

The manuscript is accompanied by a supplementary file containing notes, figures, and tables.

## Data Availability

The environmental dataset will be deposited online after formal submission. High-throughput sequences are available in the Sequence Read Archive of the National Center for Biotechnology Information (NCBI-SRA, https://ncbi.nlm.nih.gov/sra/docs/) under the BioProjects PRJNA807858, PRJNA1051030 and PRJNA1051038. The raw climatic datasets are obtained from the Climatologies at High Resolution for the Earth’s Land Surface Areas (CHELSA: https://chelsa-climate.org/).

## Author Contributions

H.L. designed this research in consultation with C.-X.H., L.-R.S., and W.-B.W.; H.L. and X.-Y.G. took the samples in the field; X.-Y.G., Z.-W.W., and H.-J.Y. conducted laboratory measurements; X.-Y.G. and H.L. conducted statistical analyses and wrote the original manuscript; and all co-authors contributed to the revision.

## References

1. Adams, R. I., Miletto, M., Taylor, J. W., & Bruns, T. D. (2013). Dispersal in microbes: fungi in indoor air are dominated by outdoor air and show dispersal limitation at short distances. The ISME Journal, 7(7), 1262–1273. doi:10.1038/ismej.2013.28

2. Albrecht, J., Peters, M. K., Becker, J. N., Behler, C., Classen, A., Ensslin, A., . . . Schleuning, M. (2021). Species richness is more important for ecosystem functioning than species turnover along an elevational gradient. Nature Ecology & Evolution, 5(12), 1582-+. doi:10.1038/s41559-021-01550-9

3. Austin, A. T., Yahdjian, L., Stark, J. M., Belnap, J., Porporato, A., Norton, U., . . . Schaeffer, S. M. (2004). Water pulses and biogeochemical cycles in arid and semiarid ecosystems. Oecologia, 141(2), 221–235. doi:10.1007/s00442-004-1519-1

4. Bell, C. W., Tissue, D. T., Loik, M. E., Wallenstein, M. D., Acosta - Martinez, V., Erickson, R. A., & Zak, J. C. (2014). Soil microbial and nutrient responses to 7years of seasonally altered precipitation in a Chihuahuan Desert grassland. Global Change Biology, 20(5), 1657–1673. doi:10.1111/gcb.12418

5. Belnap, J. (2002). Nitrogen fixation in biological soil crusts from southeast Utah, USA. Biol. Fertil. Soils, 35(2), 128–135. doi:10.1007/s00374-002-0452-x

6. Bi, J., Zhang, N. L., Liang, Y., Yang, H. J., & Ma, K. P. (2012). Interactive effects of water and nitrogen addition on soil microbial communities in a semiarid steppe. Journal of Plant Ecology, 5(3), 320–329. doi:10.1093/jpe/rtr046

7. Bowker, M. A., Maestre, F. T., Eldridge, D., Belnap, J., Castillo-Monroy, A., Escolar, C., & Soliveres, S. (2014). Biological soil crusts (biocrusts) as a model system in community, landscape and ecosystem ecology. Biodiversity and Conservation, 23(7), 1619-1637. doi:10.1007/s10531-014-0658-x

8. Bowker, M. A., Maestre, F. T., & Escolar, C. (2010). Biological crusts as a model system for examining the biodiversity–ecosystem function relationship in soils. Soil Biol. Biochem., 42(3), 405–417. doi:10.1016/j.soilbio.2009.10.025

9. Bowker, M. A., Maestre, F. T., & Mau, R. L. (2013). Diversity and Patch-Size Distributions of Biological Soil Crusts Regulate Dryland Ecosystem Multifunctionality. Ecosystems, 16, 923–933. doi:10.1007/s10021-013-9644-5)

10. Bradford, J. B., Schlaepfer, D. R., Lauenroth, W. K., & Palmquist, K. A. (2020). Robust ecological drought projections for drylands in the 21st century. Global Change Biology, 26(7), 3906–3919. doi:10.1111/gcb.15075

11. Bradford, M. A., Wood, S. A., Bardgett, R. D., Black, H. J., Bonkowski, M., Eggers, T., . . . Jones, T. H. (2014). Discontinuity in the responses of ecosystem processes and multifunctionality to altered soil community composition. Proc. Natl Acad. Sci. USA, 111(40), 14478–14483.

12. Byrnes, J. E. K., Gamfeldt, L., Isbell, F., Lefcheck, J. S., Griffin, J. N., Hector, A., . . . Freckleton, R. (2014). Investigating the relationship between biodiversity and ecosystem multifunctionality: Challenges and solutions. Methods Ecol. Evol., 5(2), 111–124. doi:10.1111/2041-210x.12143

13. Canarini, A., Schmidt, H., Fuchslueger, L., Martin, V., Herbold, C. W., Zezula, D., . . . Richter, A. (2021). Ecological memory of recurrent drought modifies soil processes via changes in soil microbial community. Nature Communications, 12(1), 14. doi:10.1038/s41467-021-25675-4

14. Chen, K., Huo, T., Zhang, Y., Guo, T., & Liang, J. (2023). Response of soil organic carbon decomposition to intensified water variability co-determined by the microbial community and aggregate changes in a temperate grassland soil of northern China. Soil Biology and Biochemistry, 176, 108875.

15. Chu, H., Sun, H., Tripathi, B. M., Adams, J. M., Huang, R., Zhang, Y., & Shi, Y. (2016). Bacterial community dissimilarity between the surface and subsurface soils equals horizontal differences over several kilometers in the western Tibetan Plateau. Environ. Microbiol., 18(5), 1523–1533. doi:10.1111/1462-2920.13236

16. Clarke, K. R. (1993). NONPARAMETRIC MULTIVARIATE ANALYSES OF CHANGES IN COMMUNITY STRUCTURE. Australian Journal of Ecology, 18(1), 117–143. doi:10.1111/j.1442-9993.1993.tb00438.x

17. Cregger, M. A., Schadt, C. W., McDowell, N. G., Pockman, W. T., & Classen, A. T. (2012). Response of the soil microbial community to changes in precipitation in a semiarid ecosystem. Applied and Environmental Microbiology, 78(24), 8587–8594.

18. Delgado-Baquerizo, M., Giaramida, L., Reich, P. B., Khachane, A. N., Hamonts, K., Edwards, C., . . . Singh, B. K. (2016). Lack of functional redundancy in the relationship between microbial diversity and ecosystem functioning. Journal of Ecology, 104(4), 936–946. doi:10.1111/1365-2745.12585

19. Delgado-Baquerizo, M., Maestre, F. T., Reich, P. B., Jeffries, T. C., Gaitan, J. J., Encinar, D., . . . Singh, B. K. (2016). Microbial diversity drives multifunctionality in terrestrial ecosystems. Nature Communications, 7, 10541.

20. Dubois, M., Gilles, K. A., Hamilton, J. K., Rebers, P. A., & Smith, F. (1956). Colorimetric method for determination of sugars and related substances. Analytical Chemistry, 28(3), 350–356. doi:10.1021/ac60111a017

21. Edgar, R. C. (2018). Updating the 97% identity threshold for 16S ribosomal RNA OTUs. Bioinformatics, 34(14), 2371–2375. doi:10.1093/bioinformatics/bty113

22. Eisenhauer, N., Hines, J., Maestre, F. T., & Rillig, M. C. (2023). Reconsidering functional redundancy in biodiversity research. npj Biodiversity, 2(1). doi:10.1038/s44185-023-00015-5

23. Evans, S., Allison, S., & Hawkes, C. (2022). Microbes, memory and moisture: Predicting microbial moisture responses and their impact on carbon cycling. Functional Ecology, 36(6), 1430–1441. doi:10.1111/1365-2435.14034

24. Evans, S. E., & Wallenstein, M. D. (2014). Climate change alters ecological strategies of soil bacteria. Ecol Lett, 17(2), 155–164. doi:10.1111/ele.12206

25. Garcia-Pichel, F. (2023). The Microbiology of Biological Soil Crusts. Annu Rev Microbiol, 77, 149–171. doi:10.1146/annurev-micro-032521-015202

26. Garland, G., Banerjee, S., Edlinger, A., Miranda Oliveira, E., Herzog, C., Wittwer, R., . . . Hector, A. (2020). A closer look at the functions behind ecosystem multifunctionality: A review. Journal of Ecology, 109(2), 600–613. doi:10.1111/1365-2745.13511

27. Gonzalez, A., Germain, R. M., Srivastava, D. S., Filotas, E., Dee, L. E., Gravel, D., . . . Loreau, M. (2020). Scaling-up biodiversity-ecosystem functioning research. Ecology Letters, 23(4), 757–776. 10.1111/ele.13456

28. Gotelli, N. J., Ulrich, W., & Maestre, F. T. (2011). Randomization tests for quantifying species importance to ecosystem function. Methods in Ecology and Evolution, 2(6), 634–642. doi:10.1111/j.2041-210X.2011.00121.x

29. Guo, X., Li, H., Huo, D., Hu, C., Li, R., Zhang, S., & Song, L. (2023). Aridity modulates biogeographic distribution and community assembly of cyanobacterial morphotypes in drylands. Fems Microbiology Ecology, 99(6). doi:10.1093/femsec/fiad053

30. Hagan, J. G., Vanschoenwinkel, B., & Gamfeldt, L. (2021). We should not necessarily expect positive relationships between biodiversity and ecosystem functioning in observational field data. Ecology Letters, 24(12), 2537–2548. 10.1111/ele.13874

31. Haiyan, D. A. I., & Haimei, W. (2021). Influence of rainfall events on soil moisture in a typical steppe of Xilingol. Physics and Chemistry of the Earth, 121, 5. doi:10.1016/j.pce.2020.102964

32. Holguin, J., Collins, S. L., & McLaren, J. R. (2022). Belowground responses to altered precipitation regimes in two semi-arid grasslands. Soil Biology & Biochemistry, 171, 12. doi:10.1016/j.soilbio.2022.108725

33. Hooper, D. U., Adair, E. C., Cardinale, B. J., Byrnes, J. E. K., Hungate, B. A., Matulich, K. L., . . . O’Connor, M. I. (2012). A global synthesis reveals biodiversity loss as a major driver of ecosystem change. Nature, 486(7401), 105–U129. doi:10.1038/nature11118

34. Hu, C. X., & Liu, Y. D. (2003). Primary succession of algal community structure in desert soil. Acta Botanica Sinica, 45(8), 917–924.

35. Hu, W. G., Ran, J. Z., Dong, L. W., Du, Q. J., Ji, M. F., Yao, S. R., . . . Deng, J. M. (2021). Aridity-driven shift in biodiversity-soil multifunctionality relationships. Nature Communications, 12(1), 15. doi:10.1038/s41467-021-25641-0

36. Huang, G., Li, Y., & Su, Y. G. (2015). Effects of increasing precipitation on soil microbial community composition and soil respiration in a temperate desert, Northwestern China. Soil Biology & Biochemistry, 83, 52–56. doi:10.1016/j.soilbio.2015.01.007

37. IPCC. (2021). In V. Masson-Delmotte, P. Zhai, A. Pirani, S. L. Connors, C. Péan, S. Berger, N. Caud, Y. Chen, L. Goldfarb, M. I. Gomis, M. Huang, K. Leitzell, E. Lonnoy, J. B. R. Matthews, T. K. Maycock, T. Waterfield, O. Yelekçi, R. Yu, & B. Zhou (Eds.), Climate Change 2021: The Physical Science Basis. Contribution of Working Group I to the Sixth Assessment Report of the Intergovernmental Panel on Climate Change (pp. 2061–2086). Cambridge, United Kingdom and New York, NY, USA: Cambridge University Press.

38. Isbell, F., Cowles, J., Dee, L. E., Loreau, M., Reich, P. B., Gonzalez, A., . . . Schmid, B. (2018). Quantifying effects of biodiversity on ecosystem functioning across times and places. Ecology Letters, 21(6), 763–778.

39. Jiao, S., Chu, H., Zhang, B., Wei, X., Chen, W., & Wei, G. (2022). Linking soil fungi to bacterial community assembly in arid ecosystems. iMeta, *1*(1 %U https://onlinelibrary.wiley.com/doi/10.1002/imt2.2).

40. Jing, X., Sanders, N. J., Shi, Y., Chu, H., Classen, A. T., Zhao, K., . . . He, J. S. (2015). The links between ecosystem multifunctionality and above- and belowground biodiversity are mediated by climate. Nat. Commun., 6, 8159. doi:10.1038/ncomms9159

41. Karp, D. S., Rominger, A. J., Zook, J., Ranganathan, J., Ehrlich, P. R., & Daily, G. C. (2012). Intensive agriculture erodes β-diversity at large scales. Ecology Letters, 15(9), 963–970. 10.1111/j.1461-0248.2012.01815.x

42. Lange, O. L., & Belnap, J. (2016). How Biological Soil Crusts Became Recognized as a Functional Unit: A Selective History. In Biological Soil Crusts: An Organizing Principle in Drylands (pp. 15-33).

43. Legendre, P. (2014). Interpreting the replacement and richness difference components of beta diversity. Global Ecology and Biogeography, 23(11), 1324–1334. doi:10.1111/geb.12207

44. Leibold, M. A., Chase, J. M., & Ernest, S. K. M. (2017). Community assembly and the functioning of ecosystems: how metacommunity processes alter ecosystems attributes. Ecology, 98(4), 909–919. 10.1002/ecy.1697

45. Lenderink, G., & Fowler, H. J. (2017). Understanding rainfall extremes. Nature Climate Change, 7(6), 391–393. doi:10.1038/nclimate3305

46. Li, H., Chen, Y., Yu, G., Rossi, F., Huo, D., De Philippis, R., . . . Li, R. (2021). Multiple diversity facets of crucial microbial groups in biological soil crusts promote soil multifunctionality. Global Ecology and Biogeography, 30(6), 1204–1217. 10.1111/geb.13295

47. Li, H., Huo, D., Wang, W., Chen, Y., Cheng, X., Yu, G., & Li, R. (2020). Multifunctionality of biocrusts is positively predicted by network topologies consistent with interspecies facilitation. Mol Ecol, 29(8), 1560–1573. doi:10.1111/mec.15424

48. Li, H., Li, R., Rossi, F., Li, D., De Philippis, R., Hu, C., & Liu, Y. (2016). Differentiation of microbial activity and functional diversity between various biocrust elements in a heterogeneous crustal community. Catena, 147, 138–145. doi:10.1016/j.catena.2016.07.008

49. Loreau, M., & Hector, A. (2001). Partitioning selection and complementarity in biodiversity experiments. Nature, 412(6842), 72–76. doi:10.1038/35083573

50. Maestre, F. T., Quero, J. L., Gotelli, N. J., Escudero, A., Ochoa, V., Delgado-Baquerizo, M., . . . Zaady, E. (2012). Plant species richness and ecosystem multifunctionality in global drylands. Science, 335(6065), 214–218. doi:10.1126/science.1215442

51. Maurer, G. E., Hallmark, A. J., Brown, R. F., Sala, O. E., & Collins, S. L. (2020). Sensitivity of primary production to precipitation across the United States. Ecology Letters, 23(3), 527–536. doi:10.1111/ele.13455

52. Mori, A. S., Isbell, F., Fujii, S., Makoto, K., Matsuoka, S., & Osono, T. (2016). Low multifunctional redundancy of soil fungal diversity at multiple scales. Ecol. Lett., 19(3), 249–259. doi:10.1111/ele.12560

53. Mori, A. S., Isbell, F., & Seidl, R. (2018). beta-Diversity, Community Assembly, and Ecosystem Functioning. Trends Ecol Evol, 33(7), 549–564. doi:10.1016/j.tree.2018.04.012

54. Mori, H., Maruyama, F., Kato, H., Toyoda, A., Dozono, A., Ohtsubo, Y., . . . Kurokawa, K. (2013). Design and Experimental Application of a Novel Non-Degenerate Universal Primer Set that Amplifies Prokaryotic 16S rRNA Genes with a Low Possibility to Amplify Eukaryotic rRNA Genes. DNA Research, 21(2), 217–227. doi:10.1093/dnares/dst052

55. Nagelkerke, N. J. D. (1991). A note on a general definition of the coefficient of determination. Biometrika, 78, 691–692.

56. Ni, Y., Yang, T., Ma, Y., Zhang, K., Soltis, P. S., Soltis, D. E., . . . Waring, B. G. (2021). Soil pH determines bacterial distribution and assembly processes in natural mountain forests of eastern China. Global Ecology and Biogeography, 30(11), 2164–2177. doi:10.1111/geb.13373

57. Nielsen, U. N., & Ball, B. A. (2015). Impacts of altered precipitation regimes on soil communities and biogeochemistry in arid and semi-arid ecosystems. Global Change Biology, 21(4), 1407–1421. doi:10.1111/gcb.12789

58. Nübel, U., Garcia-Pichel, F., & Muyzer, G. (1997). PCR primers to amplify 16S rRNA genes from cyanobacteria. Applied and Environmental Microbiology, 63(8), 3327–3332.

59. Ombadi, M. (2023). Global warming intensifies rainfall in mountainous regions. Nature, 2. doi:10.1038/d41586-023-02001-0

60. Pedros-Alio, C. (2012). The Rare Bacterial Biosphere. Annual Review of Marine Science*, Vol* 4, *4*, 449–466. doi:10.1146/annurev-marine-120710-100948

61. Rajeev, L., da Rocha, U. N., Klitgord, N., Luning, E. G., Fortney, J., Axen, S. D., . . . Mukhopadhyay, A. (2013). Dynamic cyanobacterial response to hydration and dehydration in a desert biological soil crust. Isme Journal, 7(11), 2178–2191. doi:10.1038/ismej.2013.83

62. Rath, K. M., Fierer, N., Murphy, D. V., & Rousk, J. (2019). Linking bacterial community composition to soil salinity along environmental gradients. Isme Journal, 13(3), 836–846. doi:10.1038/s41396-018-0313-8

63. Ren, X., Sha, Y., Shi, Z., & Liu, X. (2021). Response of summer extreme precipitation over East Asia during the mid-Holocene versus future global warming. Global and Planetary Change, 197. doi:10.1016/j.gloplacha.2020.103398

64. Rolls, R. J., Deane, D. C., Johnson, S. E., Heino, J., Anderson, M. J., & Ellingsen, K. E. (2023). Biotic homogenisation and differentiation as directional change in beta diversity: synthesising driver-response relationships to develop conceptual models across ecosystems. Biol Rev Camb Philos Soc, 98(4), 1388–1423. doi:10.1111/brv.12958

65. Rousk, J., Bååth, E., Brookes, P. C., Lauber, C. L., Lozupone, C., Caporaso, J. G., . . . Fierer, N. (2010). Soil bacterial and fungal communities across a pH gradient in an arable soil. Isme Journal, 4(10), 1340–1351. doi:10.1038/ismej.2010.58

66. Sasaki, T., Ishii, N. I., Makishima, D., Sutou, R., Goto, A., Kawai, Y., . . . Eisenhauer, N. (2022). Plant and microbial community composition jointly determine moorland multifunctionality. Journal of Ecology, 110(10), 2507–2521. doi:10.1111/1365-2745.13969

67. Seaton, F. M., George, P. B. L., Lebron, I., Jones, D. L., Creer, S., & Robinson, D. A. (2020). Soil textural heterogeneity impacts bacterial but not fungal diversity. Soil Biology & Biochemistry, 144. doi:10.1016/j.soilbio.2020.107766

68. Shang, H., Xu, M., Zhao, F., & Tijjani, S. B. (2019). Spatial and Temporal Variations in Precipitation Amount, Frequency, Intensity, and Persistence in China, 1973-2016. Journal of Hydrometeorology, 20(11), 2215- 2227. doi:10.1175/jhm-d-19-0032.1

69. Soliveres, S., van der Plas, F., Manning, P., Prati, D., Gossner, M. M., Renner, S. C., . . . Allan, E. (2016). Biodiversity at multiple trophic levels is needed for ecosystem multifunctionality. Nature, 536(7617), 456-+.

70. Stein, A., Gerstner, K., & Kreft, H. (2014). Environmental heterogeneity as a universal driver of species richness across taxa, biomes and spatial scales. Ecology Letters, 17(7), 866–880. 10.1111/ele.12277

71. Tilman, D., Isbell, F., & Cowles, J. M. (2014). Biodiversity and Ecosystem Functioning. In D. J. Futuyma (Ed.), Annual Review of Ecology, Evolution, and Systematics*, Vol* 45 (Vol. 45, pp. 471-493).

72. van der Plas, F., Hennecke, J., Chase, J. M., van Ruijven, J., & Barry, K. E. (2023). Universal beta-diversity- functioning relationships are neither observed nor expected. Trends Ecol Evol, 38(6), 532–544. doi:10.1016/j.tree.2023.01.008

73. van der Plas, F., Manning, P., Soliveres, S., Allan, E., Scherer-Lorenzen, M., Verheyen, K., . . . Fischer, M. (2016). Biotic homogenization can decrease landscape-scale forest multifunctionality. Proc Natl Acad Sci U S A, 113(13), 3557–3562. doi:10.1073/pnas.1517903113

74. Voriskova, J., Elberling, B., & Prieme, A. (2019). Fast response of fungal and prokaryotic communities to climate change manipulation in two contrasting tundra soils. Environmental Microbiome, 14(1), 15. doi:10.1186/s40793-019-0344-4

75. Wagg, C., Hautier, Y., Pellkofer, S., Banerjee, S., Schmid, B., & van der Heijden, M. G. (2021). Diversity and asynchrony in soil microbial communities stabilizes ecosystem functioning. Elife, 10. doi:10.7554/eLife.62813

76. Wall, D. H. (1999). Biodiversity and ecosystem functioning: A special issue devoted to belowground biodiversity in soils and freshwater and marine sediments. Bioscience, 49(2), 107–108.

77. Wang, S., Lamy, T., Hallett, L. M., & Loreau, M. (2019). Stability and synchrony across ecological hierarchies in heterogeneous metacommunities: linking theory to data. Ecography, 42(6), 1200–1211. doi:10.1111/ecog.04290

78. Wang, X., Zhang, Q., Zhang, Z., Li, W., Liu, W., Xiao, N., . . . Han, X. (2023). Decreased soil multifunctionality is associated with altered microbial network properties under precipitation reduction in a semiarid grassland. iMeta, 2(2), e106.

79. Weber, B., Belnap, J., Budel, B., Antoninka, A. J., Barger, N. N., Chaudhary, V. B., . . . Bowker, M. A. (2022). What is a biocrust? A refined, contemporary definition for a broadening research community. Biol Rev Camb Philos Soc, 97(5), 1768-1785. doi:10.1111/brv.12862

80. Winfree, R., Reilly, J. R., Bartomeus, I., Cariveau, D. P., Williams, N. M., & Gibbs, J. (2018). Species turnover promotes the importance of bee diversity for crop pollination at regional scales. Science, 359(6377), 791–793. doi:10.1126/science.aao2117

81. Wu, Q., Yue, K., Ma, Y., Hedenec, P., Cai, Y., Chen, J., . . . Li, Y. (2022). Contrasting effects of altered precipitation regimes on soil nitrogen cycling at the global scale. Global Change Biology, 28(22), 6679–6695. doi:10.1111/gcb.16392

82. Xi, N., & Bloor, J. M. G. (2016). Interactive effects of precipitation and nitrogen spatial pattern on carbon use and functional diversity in soil microbial communities. Applied Soil Ecology, 100, 207–210.

83. Xu, H. F., Raanan, H., Dai, G. Z., Oren, N., Berkowicz, S., Murik, O., . . . Qiu, B. S. (2021). Reading and surviving the harsh conditions in desert biological soil crust: the cyanobacterial viewpoint. FEMS Microbiology Reviews, 45(6), 19. doi:10.1093/femsre/fuab036

84. Yao, Y. F., Liu, J., Wang, Z., Wei, X. R., Zhu, H. S., Fu, W., & Shao, M. G. (2020). Responses of soil aggregate stability, erodibility and nutrient enrichment to simulated extreme heavy rainfall. Science of the Total Environment, 709, 9. doi:10.1016/j.scitotenv.2019.136150

85. Zhang, J. W., Feng, Y. Z., Maestre, F. T., Berdugo, M., Wang, J. T., Coleine, C., . . . Delgado-Baquerizo, M. (2023). Water availability creates global thresholds in multidimensional soil biodiversity and functions. Nature Ecology & Evolution, 7(7), 1002-+. doi:10.1038/s41559-023-02071-3

86. Zhang, W., Furtado, K., Wu, P., Zhou, T., Chadwick, R., Marzin, C., . . . Sexton, D. (2021). Increasing precipitation variability on daily-to-multiyear time scales in a warmer world. Science Advances, 7(31), eabf8021.

