## Supplemental Information for "Climate change impacts dryland multifunctionality: A cascade of precipitation variability to species replacement in multitrophic microbiota"

**Supplemental Information** contains **1** Note, **13** Figures, and **7** Tables.

**Supplementary Note 1** Additional information about the locations of study sites, precipitation parameters, regimes, edaphic property measurement, and function assessment of biocrusts.

**Supplementary Figure 1.** Trade-offs and redundancy among individual soil functions. The lower triangular matrix presents pairwise linear relationships between soil variables, with the gray areas representing the 95% confidence interval; the upper matrix shows Spearman's correlation coefficients (*r*) for each fitting, with combinations having high correlation ($|r|>0.6$) highlighted in bold; the curves on the diagonal illustrate the density distribution of variables; significant levels of the Spearman's correlation are indicated as $^{***}p < 0.001$, $^{**}p < 0.01$, and $^{*}p < 0.05$. Chl*a*: chlorophyll *a* content, WHC: water-holding capacity, EPS: exopolysaccharide, TOC: total organic carbon, S-SC: soil sucrase, S-$\beta$-GC: soil $\beta$-glucosidase, TN: total nitrogen content, $NO_3^-$: nitrate concentration, $NH_4^+$: ammonium content, TP: total phosphorus content, $PO_4^{3-}$: phosphate radical content, S-AKP: soil alkaline phosphatase.

**Supplementary Figure 2.** Relationships between pairwise climatic and environmental variables. The lower triangular matrix presents the pairwise linear relationships between variables, with the gray areas showing the 95% confidence interval; the upper matrix presents Spearman's correlation coefficients (*r*) for each fitting and highly correlated combinations ($|r| > 0.6$) highlighted in bold; the curves on the diagonal illustrate the density distribution of variables; significant levels of the Spearman's correlation coefficients are indicated as $^{***}p < 0.001$, $^{**}p < 0.01$, and $^{*}p < 0.05$. Environmental variables include geographic variables (longitude, latitude, and elevation), soil property variables (soil pH and salinity), and variables related to precipitation regimes (MAP: mean annual precipitation; Freq: mean annual precipitation frequency; Var: daily precipitation variability).

**Supplementary Figure 3.** Relationships between pairwise differences in environmental variables. The lower triangular matrix presents the pairwise linear relationships between variables and the gray areas show the 95% confidence interval; the upper matrix presents Spearman's correlation coefficients (*r*) of each fitting; the curves on the diagonal show the density distribution of variables; significant levels of the Spearman's correlation coefficients are indicated as $^{***}p < 0.001$, $^{**}p < 0.01$, and $^{*}p < 0.05$.

**Supplementary Figure 4.** Hypothesized models of effects on variance in multi-functions ($\beta$-multifunctionality) in the drylands of northwestern China. The predictors include microbial community dissimilarity (species replacement and richness differences), soil property difference (ΔpH, ΔSalinity, and ΔC:N), precipitation regime difference (ΔMAP, ΔVar, and ΔFreq) and geographic difference (ΔElevation). The serial numbers in the figure correspond to detailed references that have previously proven the path (see below).

**Supplementary Figure 5.** Contribution (**a**, **b**) and relative contribution (**c**, **d**) of species to microbial $\beta$-diversity based on SIPMER analysis. Species importance for soil multifunctionality is calculated based on the presence-absence matrix (**a**, **c**) and abundance matrix (**b**, **d**).

**Supplementary Figure 6.** History of precipitation regime across drylands of northwestern China and its impact on soil $\alpha$-multifunctionality, based on the **1-mm** effective precipitation threshold. (**a**) The annual changes of historical precipitation in four regions from 1979 to 2016, with shaded areas representing standard deviation; (**b**) The relationship between precipitation characteristics and soil $\alpha$-multifunctionality fitted by GLMs; averaging (avg.) and threshold-based $\alpha$-multifunctionality ($\alpha$-MF) at 25%, 50% and 75% were evaluated respectively; nonsignificant relationships ($p > 0.05$) are denoted by dashed lines and the estimated coefficients are plotted in the blank space. MAP: mean annual precipitation; Freq: mean annual precipitation frequency; Var: daily precipitation variability.

**Supplementary Figure 7.** Relationships between microbial $\alpha$-diversity indices and abiotic factors, based on Spearman's correlation coefficients. Only significant coefficients ($p < 0.05$) are annotated. Bac: bacteria; Cya: Cyanobacteria; Fgi: Fungi; MPD: standardized effect size of mean phylogenetic distance; MAP: mean annual precipitation; Freq: mean annual precipitation frequency; Var: daily precipitation variability.

**Supplementary Figure 8.** Spatial turnover of microbial communities along geographic distance. The gray areas represent the 95% confidence interval. The significant levels are indicated as $^{***}p < 0.001$, $^{**}p < 0.01$, and $^{*}p < 0.05$.

**Supplementary Figure 9.** Piecewise structural equation modeling of environmental difference, biotic predictors, and asynchrony-based $\beta$-multifunctionality at **25%**. The models are built for bacteria (**a**), cyanobacteria (**b**), and fungi (**c**), respectively. The connection networks represent the relationships among the differences in geographical (ΔElevation), precipitation regime (ΔMAP,

ΔFreq, and ΔVar), soil property (ΔpH, ΔSalinity, and ΔC:N), and biotic (*Repl* and $R_{\text{diff}}$) factors. Different categories of predictors are grouped into the same box for graphical simplicity. The significant levels of the path coefficient are indicated as $^{***}p < 0.001$, $^{**}p < 0.01$, and $^{*}p < 0.05$, and only significant ($p < 0.05$) and powerful ($|\lambda| > 0.2$) paths are presented; the width of arrows is proportional to the value of standard path coefficients. The global fit tests (Fisher's *C* test, *p*-value, $R^2$, AIC, and BIC) are in the upper right corner. AIC: Akaike information criterion; BIC: Bayesian information criterion. ΔMAP: mean annual precipitation; ΔFreq: annual precipitation frequency; ΔVar: daily precipitation variability; *Repl*: species replacement; $R_{\text{diff}}$: richness difference.

**Supplementary Figure 10.** Piecewise structural equation modeling of environmental difference, biotic predictors, and asynchrony-based *β*-multifunctionality at **50%**. The models are built for bacteria (**a**), cyanobacteria (**b**), and fungi (**c**), respectively. The connection networks represent the relationships among the differences in geographical (ΔElevation), precipitation regime (ΔMAP, ΔFreq, and ΔVar), soil property (ΔpH, ΔSalinity, and ΔC:N), and biotic (*Repl* and $R_{\text{diff}}$) factors. Different categories of predictors are grouped into the same box for graphical simplicity. The significant levels of the path coefficient are indicated as $^{***}p < 0.001$, $^{**}p < 0.01$, and $^{*}p < 0.05$, and only significant ($p < 0.05$) and powerful ($|\lambda| > 0.2$) paths are presented; the width of arrows is proportional to the value of standard path coefficients. The global fit tests (Fisher's *C* test, *p*-value, $R^2$, AIC, and BIC) are in the upper right corner. AIC: Akaike information criterion; BIC: Bayesian information criterion. ΔMAP: mean annual precipitation; ΔFreq: annual precipitation frequency; ΔVar: daily precipitation variability; *Repl*: species replacement; $R_{\text{diff}}$: richness difference.

**Supplementary Figure 11.** Piecewise structural equation modeling of environmental difference, biotic predictors, and asynchrony-based *β*-multifunctionality at **75%**. The models are built for bacteria (**a**), cyanobacteria (**b**), and fungi (**c**), respectively. The connection networks represent the relationships among the differences in geographical (ΔElevation), precipitation regime (ΔMAP, ΔFreq, and ΔVar), soil property (ΔpH, ΔSalinity, and ΔC:N), and biotic (*Repl* and $R_{\text{diff}}$) factors. Different categories of predictors are grouped into the same box for graphical simplicity. The significant levels of the path coefficient are indicated as $^{***}p < 0.001$, $^{**}p < 0.01$, and $^{*}p < 0.05$, and only significant ($p < 0.05$) and powerful ($|\lambda| > 0.2$) paths are presented; the width of arrows is proportional to the value of standard path coefficients. The global fit tests (Fisher's *C* test, *p*-value, $R^2$, AIC, and BIC) are in the upper right corner. AIC: Akaike information criterion; BIC: Bayesian information criterion. ΔMAP: mean annual precipitation; ΔFreq: annual precipitation frequency; ΔVar: daily precipitation variability; *Repl*: species replacement; $R_{\text{diff}}$: richness difference.

**Supplementary Figure 12.** Species importance for soil multifunctionality (averaging *α*-multifunctionality) based on the presence-absence matrix (**a**) and abundance matrix (**b**). The dashed lines represent the thresholds that are significantly correlated with multifunctionality. The red, blue, and black numbers represent the proportion of species that are positively, negatively, and non-significantly correlated with soil multifunctionality, respectively.

**Supplementary Figure 13.** Relationships between species importance for multifunctionality and soil property of species preference. The regressions show the nonlinear responses of species functional importance to species preference for pH (**a**)**,** salinity (**b**), and C:N ratio (**c**), and their respective thresholds. The red dashed lines indicate the thresholds of breakpoints identified by segment regressions; the black horizontal lines represent the significant thresholds ($|Z| = 1.96$) for the standard effect sizes of functional importance; the grey areas show the 95% confidence interval; species importance for multifunctionality is calculated on both presence-absence and abundance matrices.

**Supplementary Table 1.** The standardized loadings of environmental factors on each principal component ($n = 9$). MAP: mean annual precipitation; Freq: mean annual precipitation frequency; Var: daily precipitation variability.

**Supplementary Table 2.** Comparison between models for distance-based and asynchrony-based $\beta$-multifunctionality ($\beta$-MF at 25%, 50% and 75%) when selecting geographic distance (Geodist) or ΔMAP for analysis. When the asynchrony-based $\beta$-multifunctionality is used as the response variable, the estimated value in the generalized linear models (GLMs) represents the exponential coefficient of predictors, and the coefficient of determination is calculated as Nagelkerke's $R^2$. *D*s: total $\beta$-diversity; *Repl*: species replacement; $R_{\text{diff}}$: richness difference; ΔMAP: variance in mean annual precipitation; ΔFreq: variance in mean annual precipitation frequency; ΔVar: variance in precipitation variability.

**Supplementary Table 3**. Linkages between $\beta$-multifunctionality and changes in precipitation regime ($Z$-score). When the asynchrony-based $\beta$-multifunctionality is used as the response variable, the estimated value in GLMs represented the exponential coefficient of predictors, and the coefficient of determination is calculated as Nagelkerke's $R^2$. ΔMAP: variance in mean annual precipitation; ΔFreq: variance in annual precipitation frequency; ΔVar: variance in daily precipitation variability. Two effective precipitation thresholds at 0.1 mm and 1 mm are evaluated, respectively.

**Supplementary Table 4.** Linkages between microbial $\alpha$-diversity ($Z$-score) and soil $\alpha$-multifunctionality. Linear regression is used to check the relationship between $\alpha$-diversity and averaging $\alpha$-multifunctionality ($\alpha$-MF avg.), and GLM is used to check the relationships between $\alpha$-diversity and threshold-based $\alpha$-multifunctionality (threshold at 25%, 50%, 75%). The regression coefficients of $\alpha$-diversity from each fitting are presented below. MPD: standardized effect size of mean phylogenetic distance.

**Supplementary Table 5.** Comparison between multiple models for microbial $\beta$-diversity when selecting geographic distance (Geodist) or ΔMAP for analysis. *D*s: total $\beta$-diversity; *Repl*: species replacement; $R_{\text{diff.}}$: richness difference; ΔMAP: variance in mean annual precipitation; ΔFreq: variance in mean annual precipitation frequency; ΔVar: variance in precipitation variability.

**Supplementary Table 6.** Effects of abiotic and biotic (community composition) variables on distance-based $\beta$-multifunctionality (Euclidean) based on multiple regression. ΔMAP: variance in mean annual precipitation; ΔFreq: variance in annual precipitation frequency; ΔVar: variance in daily precipitation variability; *Repl*: species replacement; $R_{diff}$: richness difference.

**Supplementary Table 7.** Effects of abiotic and biotic (community composition) variables on asynchrony-based $\beta$-multifunctionality ($\beta$-MF) based on GLMs. ΔMAP: variance in mean annual precipitation; ΔFreq: variance in annual precipitation frequency; ΔVar: variance in daily precipitation variability; *Repl*: species replacement; $R_{diff}$: richness difference.

**Supplementary Note 1** Additional information about the locations of study sites, precipitation parameters, regimes, edaphic property measurement, and function assessment of biocrusts.

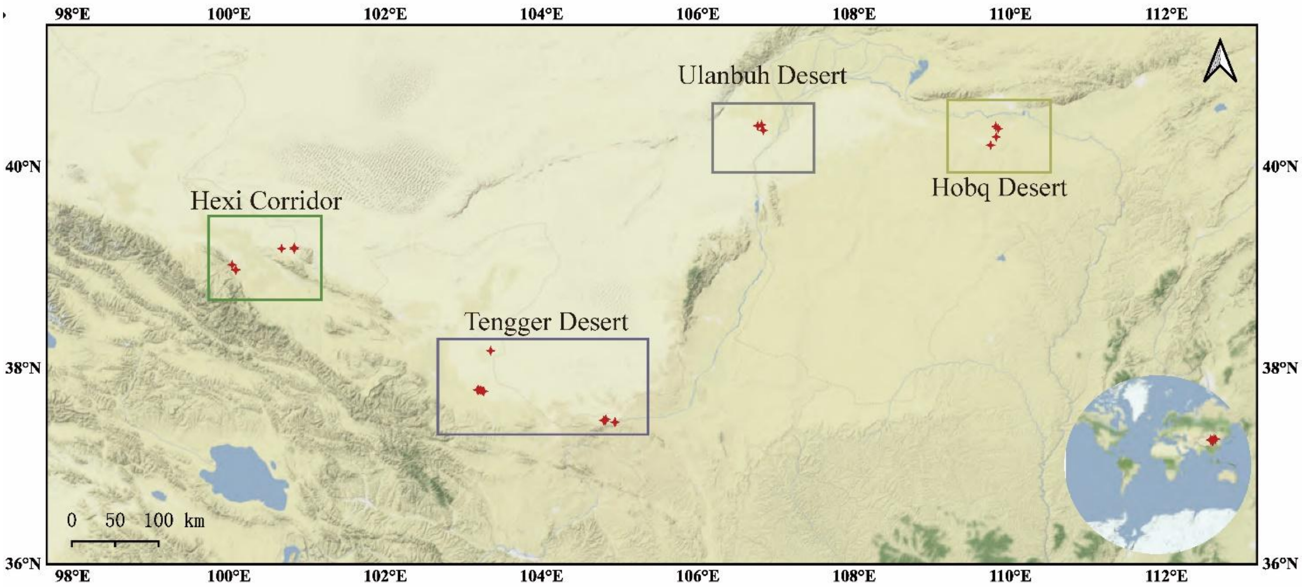


| Precipitation regimes | Sampling sites | Location |
|---|---|---|
| Regime 1 | DK1, DK2, DK3, DK4, DK5, DK6 | Ulanbuh Desert |
| Regime 2 | BTL1, BTL2, BTL3, BTL4; DLT1, DLT2; JCX1, JCX2, JCX3 | Hobq Desert |
| Regime 3 | NHZ1, NHZ2; SHW1, SHW2, SHW3, SHW4; SPT1, SPT2, SPT3, SPT4, SPT5; ZY1, ZY2, ZY3, ZY4, ZY5, ZY6, ZY7, ZY8 | Hexi Corridor<br>Tengger Desert |
| Regime 4 | DX1, DX2, DX3, DX4; GTX1, GTX2; HHT1, HHT2, HHT3, HHT4, HHT5 | Hexi Corridor<br>Tengger Desert |

**Site acronyms**: DK, *Dengkou*; BTL, *Baituliang*; DLT, *Dalate*; JCX, *Jiechaixian*; NHZ, *Nanhuzhen*; SHW, *Shahaowan*; SPT, *Shapotou*; ZY, *Zhangye*; DX, *Danxia Geopark*; GTX, *Gaotaixian*; HHT, *Huanghuatan*.

Minor rainfall events may not effectively replenish soil moisture or significantly impact soil moisture characteristics due to their inability to reach the soil surface (Groisman & Knight, 2008). Previous studies have used a range of precipitation thresholds for a ‘dry day’, from 0.1 to 10.0 mm per day (Liu, Liu, Henderson, Xu, & Zhou, 2015). The 1.0-mm threshold is most commonly used (McCabe, Legates, & Lins, 2010; McErlich et al., 2023), while standard rain gauges at Chinese meteorological stations have measured precipitation with a resolution of 0.1 mm per day for decades. Consequently, we used 0.1 mm and 1 mm as the thresholds for effective precipitation in our analysis of daily precipitation characteristics.

Samples were ground and sieved, then mixed with distilled water at a 1:5 ratio. After thorough centrifugation, we extracted the supernatant and pretreated it with a C18 column (IC Guard C18 column, CNW Technologies GmbH, Düsseldorf, Germany). The filtered portion was then analyzed using ion chromatography (Dionex™ ICS-5000+, Thermo Fisher Scientific, USA). Cation and anion concentrations were calibrated using Dionex Six Cation Standard II ($K^+$, $Ca^{2+}$, $Na^+$, $Mg^{2+}$, and $NH_4^+$) and Dionex Seven Anion Standard II ($Cl^-$, $SO_4^{2-}$, $NO_3^-$, $NO_2^-$, and $PO_4^{3-}$), respectively. Finally, soil salinity ($\mu mol \cdot g^{-1}$) was calculated as the sum of the concentrations of the 10 ions.

The content of extracellular polysaccharides (EPSs, $\mu g \cdot g^{-1}$) comprises both colloidal-EPSs (C-EPSs) and tightly bound EPSs (TB-EPSs). The extraction method adheres to the procedure described in a previous study (Chen et al., 2014). C-EPSs, being loosely bound, are easily released into the surrounding medium. Dry samples were subjected to three rounds of extraction with distilled water for 15 min in a shaker set at 30 °C and a rotational speed of 100 rpm. After thorough centrifugation, the supernatants containing C-EPSs were collected and stored for analysis. The resulting pellets were then suspended in 0.1 M $Na_2EDTA$ and shaken for 16 h at room temperature. The extracts were clarified by centrifugation, and the supernatants containing TB-EPSs were then collected and stored for subsequent analysis. We quantified the contents of C-EPSs and TB-EPSs in the non-dialyzed extracts using the phenol sulfuric acid assay (Dubois, Gilles, Hamilton, Rebers, & Smith, 1956), with glucose as the standard.

### References

Chen, L., Rossi, F., Deng, S., Liu, Y., Wang, G., Adessi, A., & De Philippis, R. (2014). Macromolecular and chemical features of the excreted extracellular polysaccharides in induced biological soil crusts of different ages. *Soil Biology & Biochemistry, 78*, 1-9.

Dubois, M., Gilles, K. A., Hamilton, J. K., Rebers, P. A., & Smith, F. (1956). **Colorimetric method for determination of sugars and related substances**. *Analytical Chemistry, 28*(3), 350-356. doi:10.1021/ac60111a017

Groisman, P. Y., & Knight, R. W. (2008). Prolonged dry episodes over the conterminous United States: New tendencies emerging during the last 40 years. *Journal of Climate, 21*(9), 1850-1862. doi:10.1175/2007jcli2013.1

Liu, X. D., Liu, B. H., Henderson, M., Xu, M., & Zhou, D. W. (2015). Observed changes in dry day frequency and prolonged dry episodes in Northeast China. *International Journal of Climatology, 35*(2), 196-214. doi:10.1002/joc.3972

McCabe, G. J., Legates, D. R., & Lins, H. F. (2010). Variability and trends in dry day frequency and dry event length in the southwestern United States. *Journal of Geophysical Research-Atmospheres, 115*, 8. doi:10.1029/2009jd012866

McErlich, C., McDonald, A., Schuddeboom, A., Vishwanathan, G., Renwick, J., & Rana, S. (2023). Positive correlation between wet-day frequency and intensity linked to universal precipitation drivers. *Nat Geosci*, 17. doi:10.1038/s41561-023-01177-4

**Supplementary Figure 1.** Trade-offs and redundancy among individual soil functions. The lower triangular matrix presents pairwise linear relationships between soil variables, with the gray areas representing the 95% confidence interval; the upper matrix shows Spearman's correlation coefficients (*r*) for each fitting, with combinations having high correlation ($|r|>0.6$) highlighted in bold; the curves on the diagonal illustrate the density distribution of variables; significant levels of the Spearman's correlation are indicated as $^{***}p < 0.001$, $^{**}p < 0.01$, and $^{*}p < 0.05$. Chl*a*: chlorophyll *a* content, WHC: water-holding capacity, EPS: exopolysaccharide, TOC: total organic carbon, S-SC: soil sucrase, S-*β*-GC: soil *β*-glucosidase, TN: total nitrogen content, $NO_3^-$: nitrate concentration, $NH_4^+$: ammonium content, TP: total phosphorus content, $PO_4^{3-}$: phosphate radical content, S-AKP: soil alkaline phosphatase.

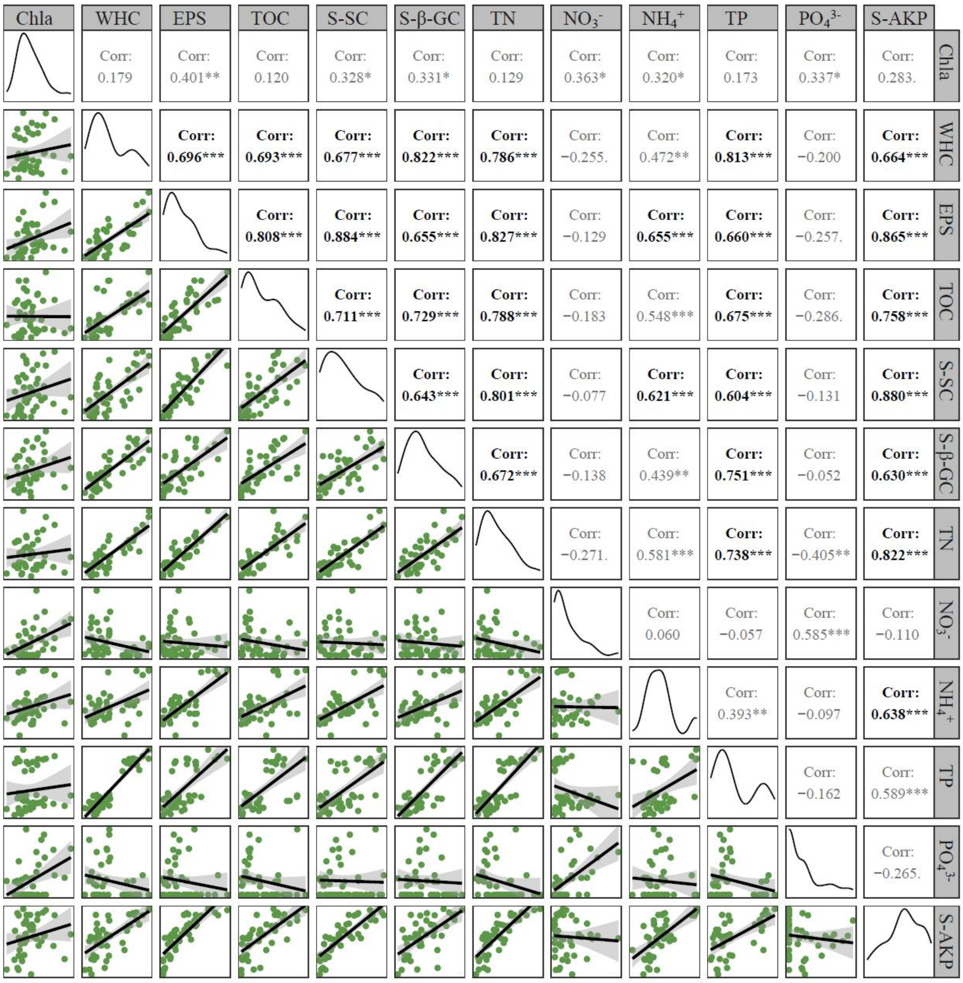

**Supplementary Figure 2.** Relationships between pairwise climatic and environmental variables. The lower triangular matrix presents the pairwise linear relationships between variables, with the gray areas showing the 95% confidence interval; the upper matrix presents Spearman's correlation coefficients ($r$) for each fitting and highly correlated combinations ($|r| > 0.6$) highlighted in bold; the curves on the diagonal illustrate the density distribution of variables; significant levels of the Spearman's correlation coefficients are indicated as $^{***}p < 0.001$, $^{**}p < 0.01$, and $^{*}p < 0.05$. Environmental variables include geographic variables (longitude, latitude, and elevation), soil property variables (soil pH and salinity), and variables related to precipitation regimes (MAP: mean annual precipitation; Freq: mean annual precipitation frequency; Var: daily precipitation variability).

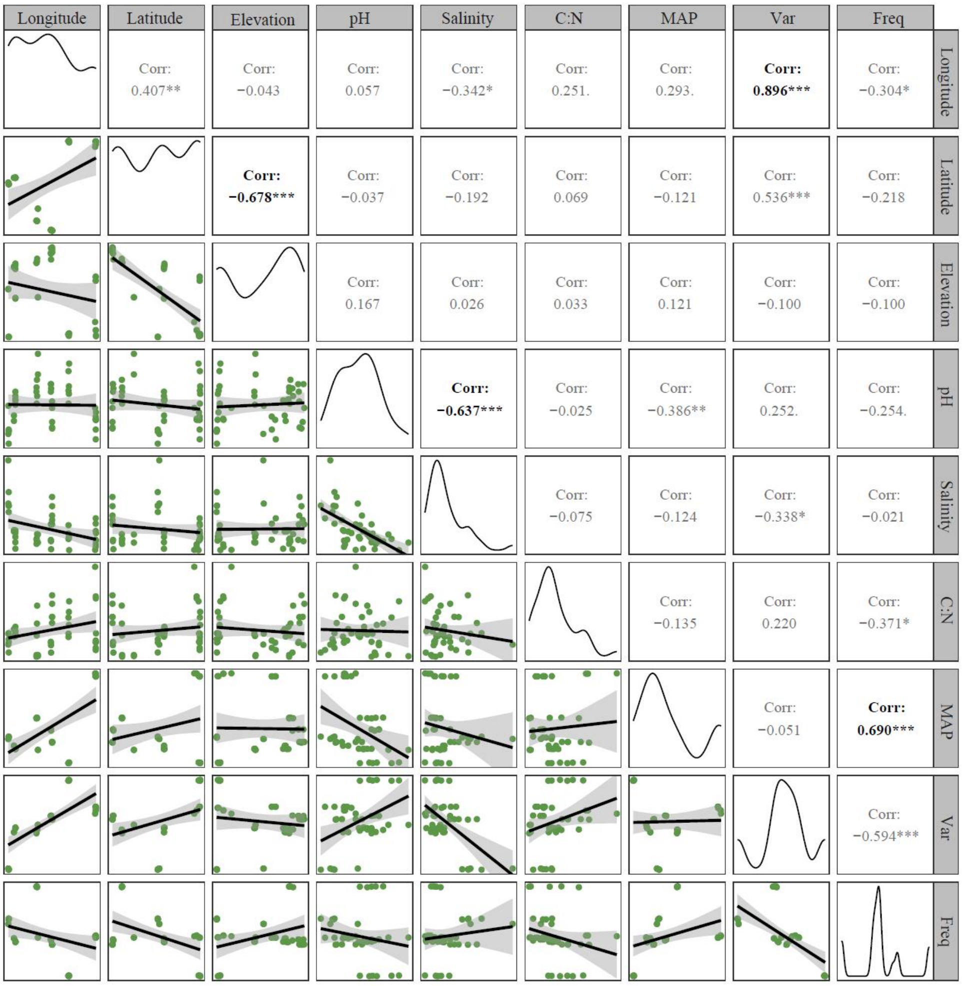

**Supplementary Figure 3.** Relationships between pairwise differences in environmental variables. The lower triangular matrix presents the pairwise linear relationships between variables and the gray areas show the 95% confidence interval; the upper matrix presents Spearman's correlation coefficients ($r$) of each fitting; the curves on the diagonal show the density distribution of variables; significant levels of the Spearman's correlation coefficients are indicated as $^{***}p < 0.001$, $^{**}p < 0.01$, and $^{*}p < 0.05$.

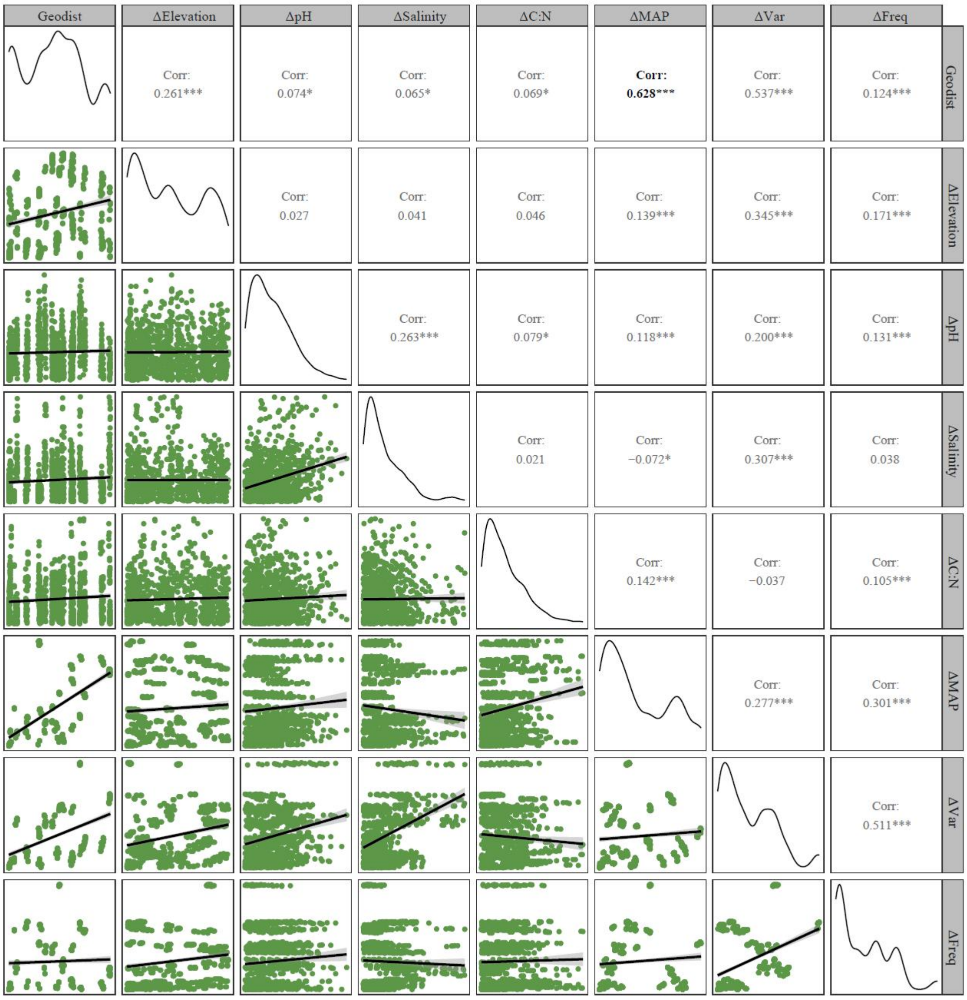

**Supplementary Figure 4.** Hypothesized models of effects on variance in multi-functions (*β*-multifunctionality) in the drylands of northwestern China. The predictors include microbial community dissimilarity (species replacement and richness differences), soil property difference (ΔpH, ΔSalinity, and ΔC:N), precipitation regime difference (ΔMAP, ΔVar, and ΔFreq) and geographic difference (ΔElevation). The serial numbers in the figure correspond to detailed references that have previously proven the path (see below).

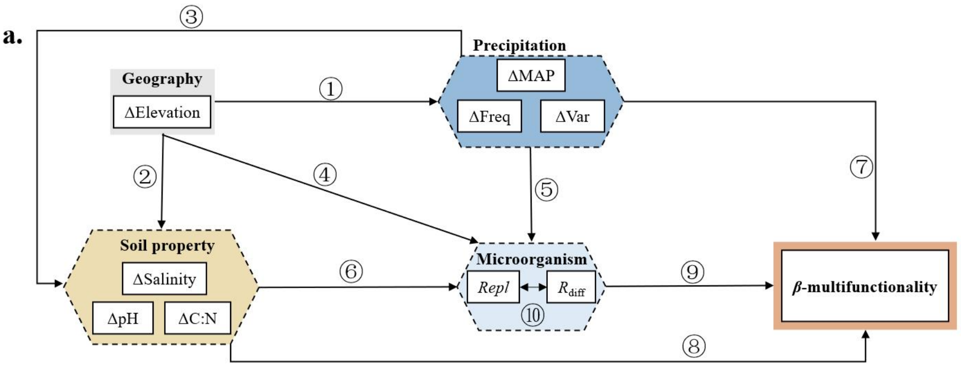


**Path：** ①②④⑤

Liu, Z.H., Fang, J., Song, B., Yang, Y., Yu, Z., Hu, J.L. et al. (2023) Stochastic processes dominate soil arbuscular mycorrhizal fungal community assembly along an elevation gradient in central Japan. *Sci Total Environ* 855: 11.

**Path：** ②

Wang, X.S., Michalet, R., He, S., and Wang, X.T. (2023) The subalpine shrub Dasiphora fruticosa alters seasonal and elevational effects on soil microbial diversity and ecosystem functions on the Tibetan Plateau. *J Appl Ecol* 60: 52-63.

**Path：** ③⑤⑥⑦⑧

Jing, X., Prager, C.M., Chen, L.T., Chu, H.Y., Gotelli, N.J., He, J.S. et al. (2022) The influence of aboveground and belowground species composition on spatial turnover in nutrient pools in alpine grasslands. *Global Ecol Biogeogr* 31: 486-500.

**Path：** ④

Gong, S., Feng, B., Jian, S.P., Wang, G.S., Ge, Z.W., and Yang, Z.L. (2022) Elevation Matters More than Season in Shaping the Heterogeneity of Soil and Root Associated Ectomycorrhizal Fungal Community. *Microbiology Spectrum* 10: 17.

**Path：** ⑧

Martinez-Almoyna, C., Thuiller, W., Chalmandrier, L., Ohlmann, M., Foulquier, A., Clement, J.C. et al. (2019) Multi-trophic beta-diversity mediates the effect of environmental gradients on the turnover of multiple ecosystem functions. *Funct Ecol* 33: 2053-2064.

**Path：** ⑨⑩

Albrecht, J., Peters, M.K., Becker, J.N., Behler, C., Classen, A., Ensslin, A. et al. (2021) Species richness is more important for ecosystem functioning than species turnover along an elevational gradient. *Nature Ecology & Evolution* 5: 1582-+.

**Supplementary Figure 5.** Contribution (**a**, **b**) and relative contribution (**c**, **d**) of species to microbial $\beta$-diversity based on SIPMER analysis. Species importance for soil multifunctionality is calculated based on the presence-absence matrix (**a**, **c**) and abundance matrix (**b**, **d**).

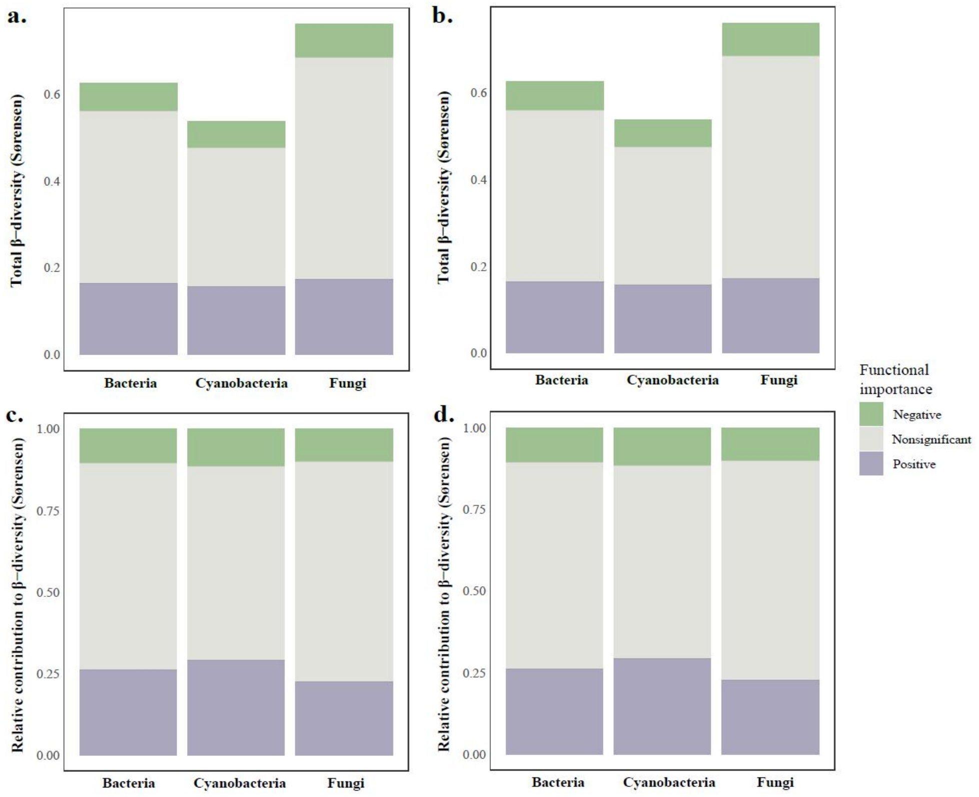

**Supplementary Figure 6.** History of precipitation regime across drylands of northwestern China and its impact on soil $\alpha$-multifunctionality, based on the **1-mm** effective precipitation threshold. (**a**) The annual changes of historical precipitation in four regions from 1979 to 2016, with shaded areas representing standard deviation; (**b**) The relationship between precipitation characteristics and soil $\alpha$-multifunctionality fitted by GLMs; averaging (avg.) and threshold-based $\alpha$-multifunctionality ($\alpha$-MF) at 25%, 50% and 75% were evaluated respectively; nonsignificant relationships ($p > 0.05$) are denoted by dashed lines and the estimated coefficients are plotted in the blank space. MAP: mean annual precipitation; Freq: mean annual precipitation frequency; Var: daily precipitation variability.

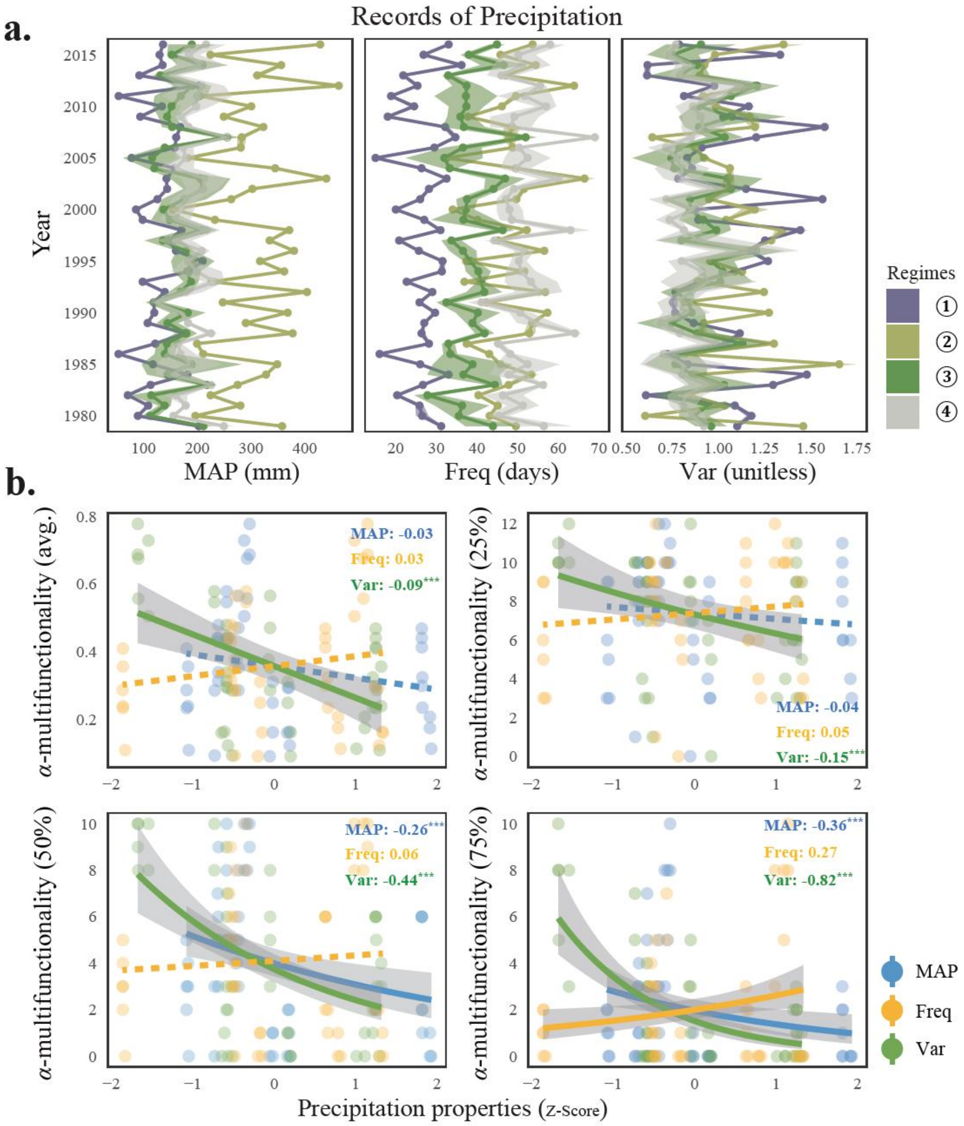

**Supplementary Figure 7.** Relationships between microbial *α*-diversity indices and abiotic factors, based on Spearman's correlation coefficients. Only significant coefficients ($p < 0.05$) are annotated. Bac: bacteria; Cya: Cyanobacteria; Fgi: Fungi; MPD: standardized effect size of mean phylogenetic distance; MAP: mean annual precipitation; Freq: mean annual precipitation frequency; Var: daily precipitation variability.

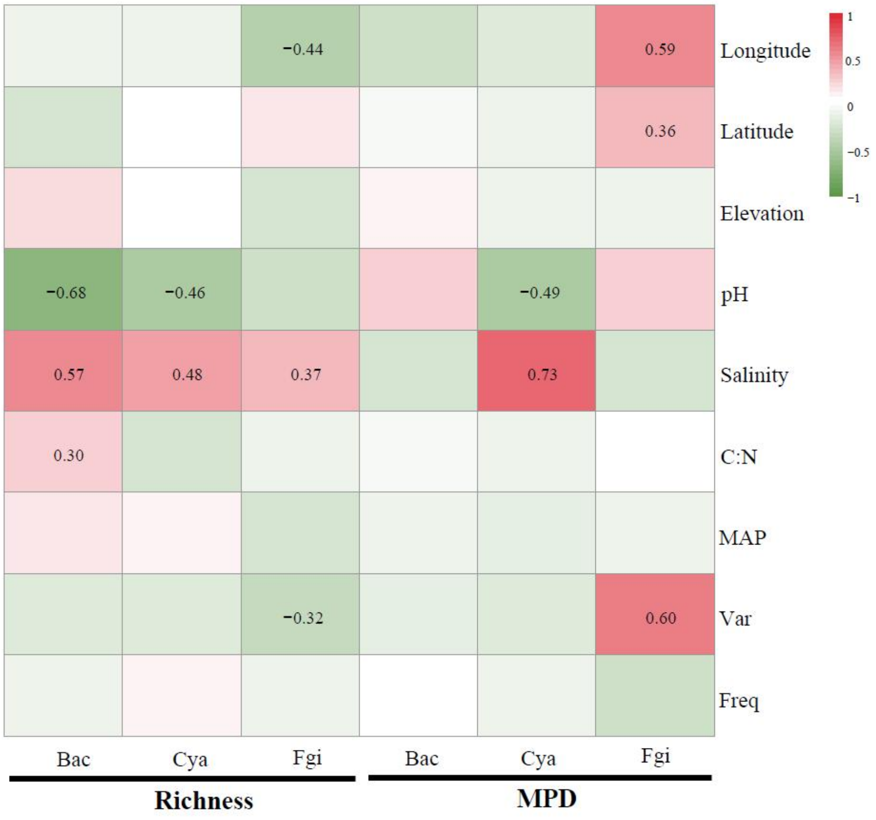

**Supplementary Figure 8.** Spatial turnover of microbial communities along geographic distance. The gray areas represent the 95% confidence interval. The significant levels are indicated as $^{***}p <$ 0.001, $^{**}p < 0.01$, and $^{*}p < 0.05$.

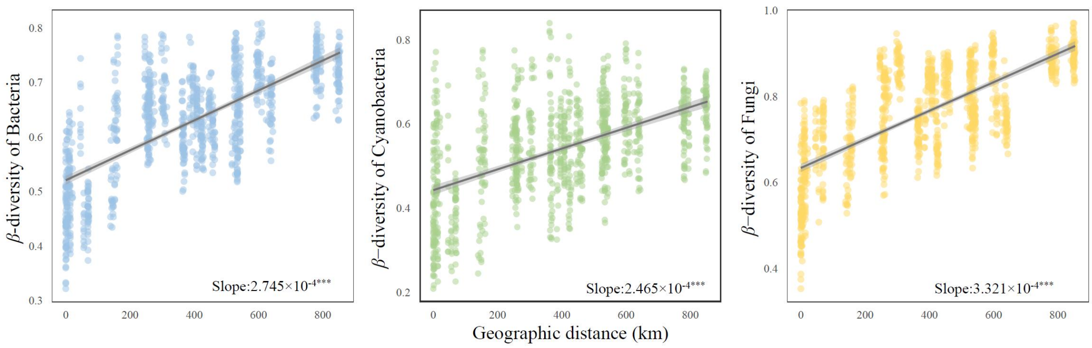

**Supplementary Figure 9.** Piecewise structural equation modeling of environmental difference, biotic predictors, and asynchrony-based $\beta$-multifunctionality at **25%**. The models are built for bacteria (**a**), cyanobacteria (**b**), and fungi (**c**), respectively. The connection networks represent the relationships among the differences in geographical (ΔElevation), precipitation regime (ΔMAP, ΔFreq, and ΔVar), soil property (ΔpH, ΔSalinity, and ΔC:N), and biotic (*Repl* and $R_{\text{diff}}$) factors. Different categories of predictors are grouped into the same box for graphical simplicity. The significant levels of the path coefficient are indicated as $^{***}p < 0.001$, $^{**}p < 0.01$, and $^{*}p < 0.05$, and only significant ($p < 0.05$) and powerful ($|\lambda| > 0.2$) paths are presented; the width of arrows is proportional to the value of standard path coefficients. The global fit tests (Fisher's $C$ test, $p$-value, $R^2$, AIC, and BIC) are in the upper right corner. AIC: Akaike information criterion; BIC: Bayesian information criterion. ΔMAP: mean annual precipitation; ΔFreq: annual precipitation frequency; ΔVar: daily precipitation variability; *Repl*: species replacement; $R_{\text{diff}}$: richness difference.

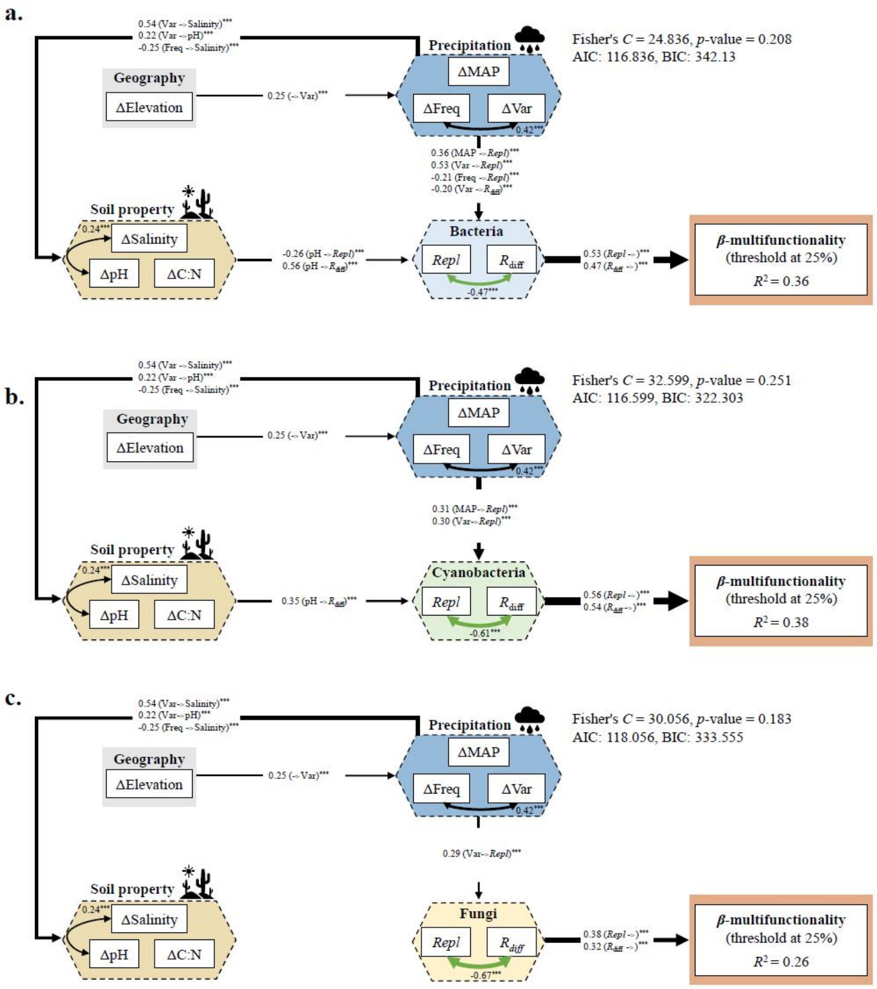

**Supplementary Figure 10.** Piecewise structural equation modeling of environmental difference, biotic predictors, and asynchrony-based $\beta$-multifunctionality at **50%**. The models are built for bacteria (**a**), cyanobacteria (**b**), and fungi (**c**), respectively. The connection networks represent the relationships among the differences in geographical (ΔElevation), precipitation regime (ΔMAP, ΔFreq, and ΔVar), soil property (ΔpH, ΔSalinity, and ΔC:N), and biotic (*Repl* and $R_{\text{diff}}$) factors. Different categories of predictors are grouped into the same box for graphical simplicity. The significant levels of the path coefficient are indicated as $^{***}p < 0.001$, $^{**}p < 0.01$, and $^{*}p < 0.05$, and only significant ($p < 0.05$) and powerful ($|\lambda| > 0.2$) paths are presented; the width of arrows is proportional to the value of standard path coefficients. The global fit tests (Fisher's $C$ test, $p$-value, $R^2$, AIC, and BIC) are in the upper right corner. AIC: Akaike information criterion; BIC: Bayesian information criterion. ΔMAP: mean annual precipitation; ΔFreq: annual precipitation frequency; ΔVar: daily precipitation variability; *Repl*: species replacement; $R_{\text{diff}}$: richness difference.

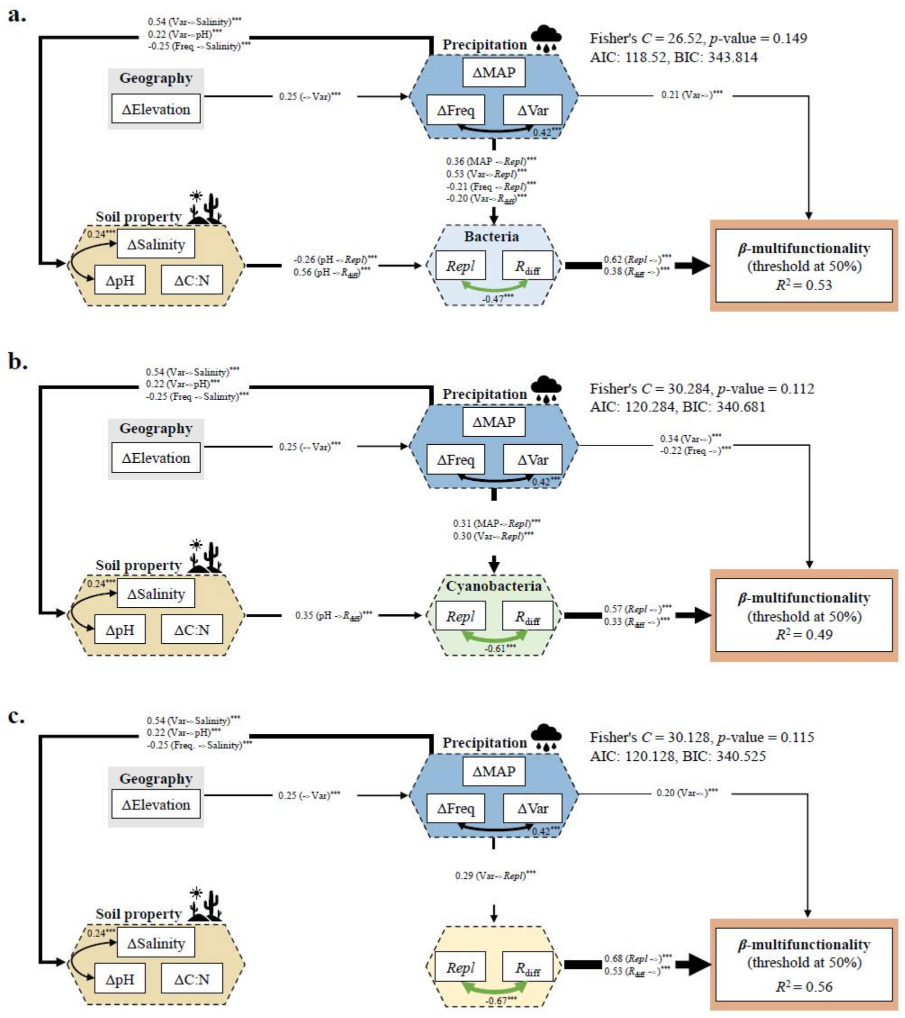

**Supplementary Figure 11.** Piecewise structural equation modeling of environmental difference, biotic predictors, and asynchrony-based $\beta$-multifunctionality at **75%**. The models are built for bacteria (**a**), cyanobacteria (**b**), and fungi (**c**), respectively. The connection networks represent the relationships among the differences in geographical (ΔElevation), precipitation regime (ΔMAP, ΔFreq, and ΔVar), soil property (ΔpH, ΔSalinity, and ΔC:N), and biotic (*Repl* and $R_{\mathrm{diff}}$) factors. Different categories of predictors are grouped into the same box for graphical simplicity. The significant levels of the path coefficient are indicated as $^{***}p < 0.001$, $^{**}p < 0.01$, and $^{*}p < 0.05$, and only significant ($p < 0.05$) and powerful ($|\lambda| > 0.2$) paths are presented; the width of arrows is proportional to the value of standard path coefficients. The global fit tests (Fisher's $C$ test, $p$-value, $R^2$, AIC, and BIC) are in the upper right corner. AIC: Akaike information criterion; BIC: Bayesian information criterion. ΔMAP: mean annual precipitation; ΔFreq: annual precipitation frequency; ΔVar: daily precipitation variability; *Repl*: species replacement; $R_{\mathrm{diff}}$: richness difference.

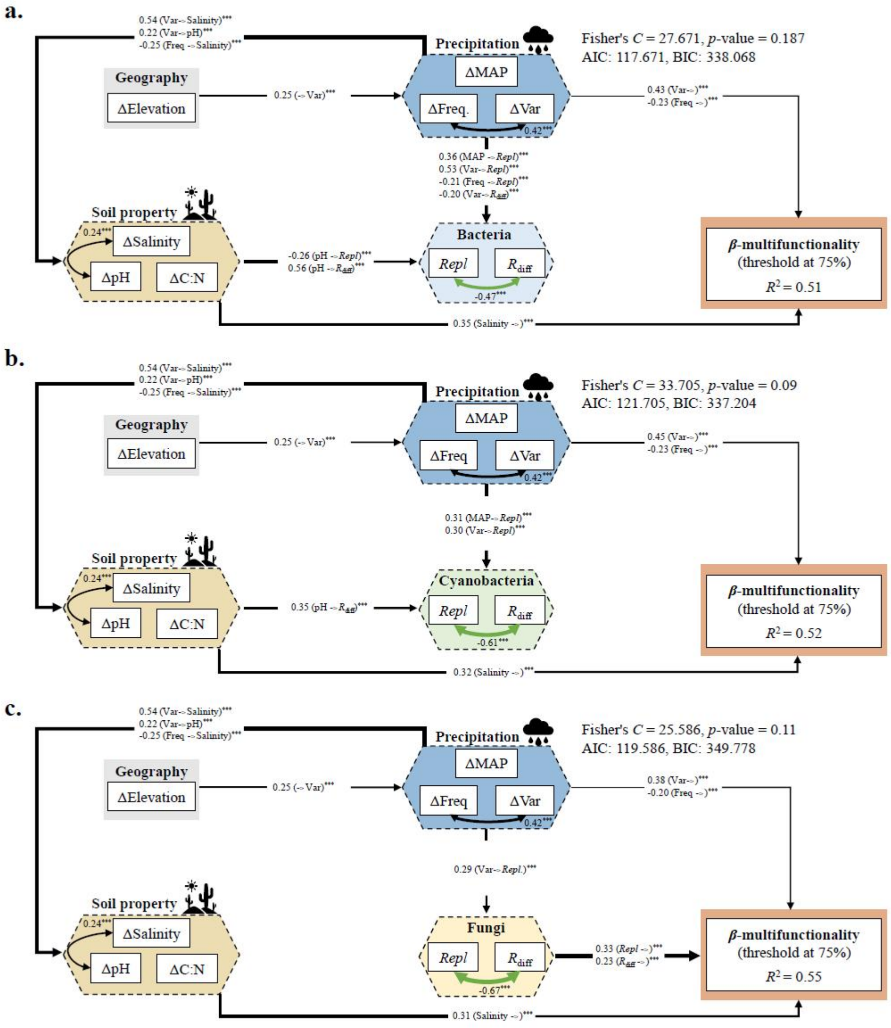

**Supplementary Figure 12.** Species importance for soil multifunctionality (averaging $\alpha$-multifunctionality) based on the presence-absence matrix (**a**) and abundance matrix (**b**). The dashed lines represent the thresholds that are significantly correlated with multifunctionality. The red, blue, and black numbers represent the proportion of species that are positively, negatively, and non-significantly correlated with soil multifunctionality, respectively.

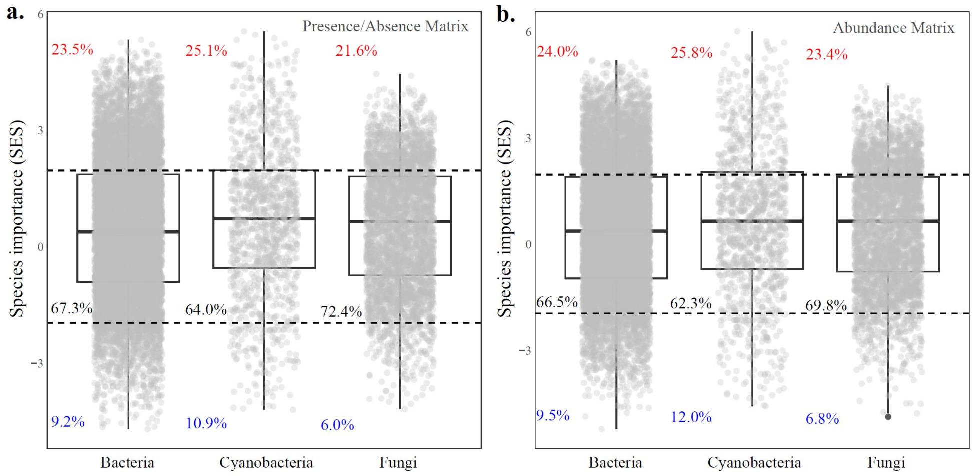

**Supplementary Figure 13.** Relationships between species importance for multifunctionality and soil property of species preference. The regressions show the nonlinear responses of species functional importance to species preference for pH (**a**), salinity (**b**), and C:N ratio (**c**), and their respective thresholds. The red dashed lines indicate the thresholds of breakpoints identified by segment regressions; the black horizontal lines represent the significant thresholds ($|Z|$ = 1.96) for the standard effect sizes of functional importance; the grey areas show the 95% confidence interval; species importance for multifunctionality is calculated on both presence-absence and abundance matrices.

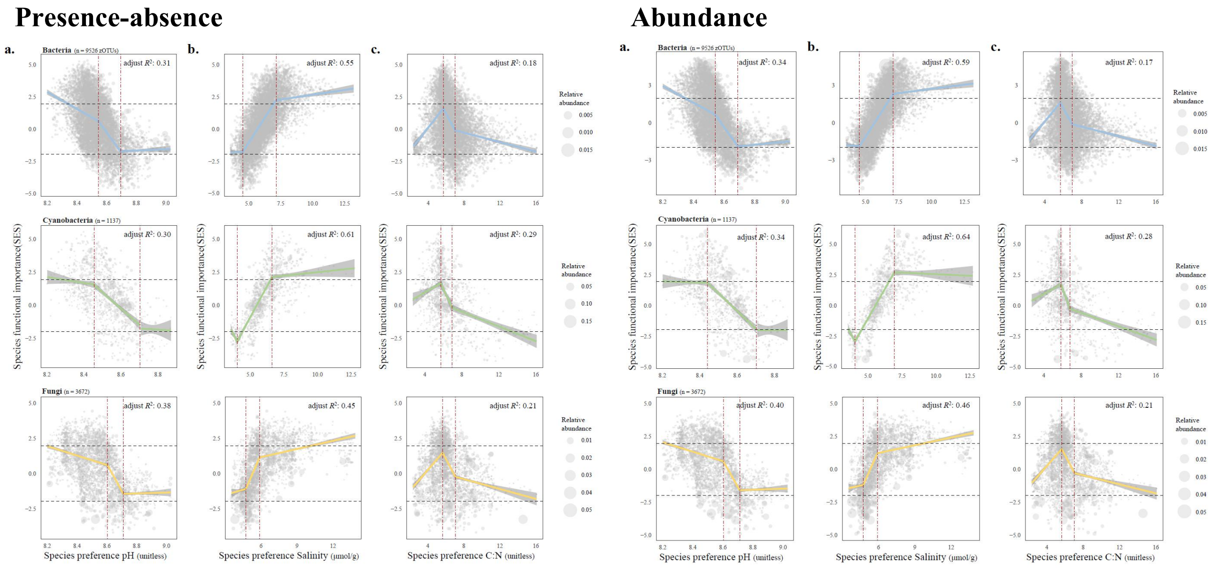

**Supplementary Table 1.** The standardized loadings of environmental factors on each principal component (*n* = 9). MAP: mean annual precipitation; Freq: mean annual precipitation frequency; Var: daily precipitation variability.

| **Factors** | RC1 | RC2 | RC3 | RC5 | RC4 |
|---|---|---|---|---|---|
| Longitude | **0.553** | **0.757** | 0.160 | -0.144 | 0.151 |
| Latitude | 0.341 | 0.285 | -0.025 | **-0.837** | 0.019 |
| Elevation | -0.055 | 0.042 | 0.008 | **0.958** | -0.066 |
| MAP | -0.186 | **0.966** | -0.070 | -0.071 | 0.025 |
| Var | **0.818** | 0.236 | 0.461 | -0.080 | 0.134 |
| Freq | **-0.880** | 0.256 | -0.011 | 0.303 | -0.114 |
| pH | 0.155 | -0.335 | **0.871** | 0.115 | -0.058 |
| Salinity | -0.108 | -0.263 | **-0.900** | 0.063 | -0.089 |
| C:N | 0.147 | 0.074 | 0.032 | -0.064 | **0.983** |
| **SS loadings** | 1.959 | 1.897 | 1.812 | 1.765 | 1.036 |
| **Proportion Var.** | 0.218 | 0.211 | 0.201 | 0.196 | 0.115 |
| **Cumulative Var.** | 0.218 | 0.428 | 0.63 | 0.826 | **0.941** |

**Supplementary Table 2.** Comparison between models for distance-based and asynchrony-based $\beta$-multifunctionality ($\beta$-MF at 25%, 50% and 75%) when selecting geographic distance (Geodist) or ΔMAP for analysis. When the asynchrony-based $\beta$-multifunctionality is used as the response variable, the estimated value in the generalized linear models (GLMs) represents the exponential coefficient of predictors, and the coefficient of determination is calculated as Nagelkerke's $R^2$. *D*s: total $\beta$-diversity; *Repl*: species replacement; $R_{diff}$: richness difference; ΔMAP: variance in mean annual precipitation; ΔFreq: variance in mean annual precipitation frequency; ΔVar: variance in precipitation variability.

| **Formula** | Adjusted $R^2$/ Nagelkerke's $R^2$ | *p*-values | AICs |
|---|---|---|---|
| **Bacteria** | | | |
| distance-based $\beta$-MF (Fit1) | 0.632 | < 0.001 | 1832.81 |
| distance-based $\beta$-MF (Fit2) | 0.618 | < 0.001 | 1869.06 |
| $\beta$-MF at 25% (Fit1) | 0.515 | < 0.001 | 4276.67 |
| $\beta$-MF at 25% (Fit2) | 0.516 | < 0.001 | 4275.54 |
| $\beta$-MF at 50% (Fit1) | 0.664 | < 0.001 | 4269.65 |
| $\beta$-MF at 50% (Fit2) | 0.658 | < 0.001 | 4281.67 |
| $\beta$-MF at 75% (Fit1) | 0.723 | < 0.001 | 4107.15 |
| $\beta$-MF at 75% (Fit2) | 0.717 | < 0.001 | 4121.80 |
| **Cyanobacteria** | | | |
| distance-based $\beta$-MF(Fit1) | 0.629 | < 0.001 | 1832.81 |
| distance-based $\beta$-MF (Fit2) | 0.621 | < 0.001 | 1860.07 |
| $\beta$-MF at 25% (Fit1) | 0.520 | < 0.001 | 4270.19 |
| $\beta$-MF at 25% (Fit2) | 0.530 | < 0.001 | 4257.43 |
| $\beta$-MF at 50% (Fit1) | 0.630 | < 0.001 | 4332.01 |
| $\beta$-MF at 50% (Fit2) | 0.641 | < 0.001 | 4311.56 |
| $\beta$-MF at 75% (Fit1) | 0.735 | < 0.001 | 4071.49 |
| $\beta$-MF at 75% (Fit2) | 0.726 | < 0.001 | 4096.05 |
| **Fungi** | | | |
| distance-based $\beta$-MF(Fit1) | 0.648 | < 0.001 | 1789.17 |
| distance-based $\beta$-MF (Fit2) | 0.643 | < 0.001 | 1801.92 |
| $\beta$-MF at 25% (Fit1) | 0.394 | < 0.001 | 4418.33 |
| $\beta$-MF at 25% (Fit2) | 0.423 | < 0.001 | 4385.75 |
| $\beta$-MF at 50% (Fit1) | 0.701 | < 0.001 | 4197.17 |
| $\beta$-MF at 50% (Fit2) | 0.702 | < 0.001 | 4195.03 |
| $\beta$-MF at 75% (Fit1) | 0.761 | < 0.001 | 3989.79 |
| $\beta$-MF at 75% (Fit2) | 0.755 | < 0.001 | 4009.00 |

Fit1: $\beta$-MF ~ ΔElevation+ΔMAP+ΔVar+ΔFreq+ΔpH+ΔSalinity+ΔC:N+*Repl*+$R_{diff}$

Fit2: $\beta$-MF ~ Geodist+ΔElevation+ΔVar+ΔFreq+ΔpH+ΔSalinity+ΔC:N+*Repl*+$R_{diff}$

**Supplementary Table 3**. Linkages between *β*-multifunctionality (*β*-MF) and changes in precipitation regime (*Z*-score). When the asynchrony-based *β*-multifunctionality is used as the response variable, the estimated value in GLMs represented the exponential coefficient of predictors, and the coefficient of determination is calculated as Nagelkerke's $R^2$. ΔMAP: variance in mean annual precipitation; ΔFreq: variance in annual precipitation frequency; ΔVar: variance in daily precipitation variability. Two effective precipitation thresholds at 0.1 mm and 1 mm are evaluated, respectively.

| | | **0.1-mm** | | | **1-mm** | | |
|---|---|---|---|---|---|---|---|
| | | ΔMAP | ΔFreq | ΔVar | ΔMAP | ΔFreq | ΔVar |
| *β*-MF (distance-based) | Estimate | n.s. | n.s. | $0.26^{***}$ | n.s. | $0.04^{*}$ | $0.26^{***}$ |
| | Adjusted $R^2$ | - | - | **0.30** | - | 0.01 | **0.29** |
| *β*-MF (threshold at 25%) | Estimate | $0.11^{***}$ | $0.07^{***}$ | $0.10^{***}$ | $0.10^{***}$ | $0.07^{***}$ | $0.15^{***}$ |
| | Nagelkerke's $R^2$ | 0.08 | 0.04 | **0.07** | 0.08 | 0.04 | **0.16** |
| *β*-MF (threshold at 50%) | Estimate | $0.05^{***}$ | $-0.03^{*}$ | $0.22^{***}$ | $0.05^{***}$ | n.s. | $0.26^{***}$ |
| | Nagelkerke's $R^2$ | 0.01 | 0.01 | **0.30** | 0.01 | - | **0.37** |
| *β*-MF (threshold at 75%) | Estimate | $-0.08^{***}$ | $-0.04^{*}$ | $0.39^{***}$ | $-0.08^{***}$ | n.s. | $0.37^{***}$ |
| | Nagelkerke's $R^2$ | 0.02 | 0.01 | **0.50** | 0.02 | - | **0.41** |

The significant levels are indicated as $^{***}$ $p < 0.001$, $^{**}$ $p < 0.01$, $^{*}$ $p < 0.05$.

**Supplementary Table 4.** Linkages between microbial $\alpha$-diversity ($Z$-score) and soil $\alpha$-multifunctionality. Linear regression is used to check the relationship between $\alpha$-diversity and averaging $\alpha$-multifunctionality ($\alpha$-MF avg.), and GLM is used to check the relationships between $\alpha$-diversity and threshold-based $\alpha$-multifunctionality (threshold at 25%, 50%, 75%). The regression coefficients of $\alpha$-diversity from each fitting are presented below. MPD: standardized effect size of mean phylogenetic distance.

| **Without considering abiotic variables** (Fit: $\alpha$-MF ~ $\alpha$-diversity)**:** | | | | |
|---|---|---|---|---|
| Predictors | $\alpha$-MF (avg.) | $\alpha$-MF (25%) | $\alpha$-MF (50%) | $\alpha$-MF (75%) |
| **Bacteria** | | | | |
| Richness | $0.097^{***}$ | $0.254^{***}$ | $0.494^{***}$ | $0.704^{***}$ |
| MPD | n.s. | n.s. | n.s. | n.s. |
| **Cyanobacteria** | | | | |
| Richness | $0.078^{**}$ | $0.225^{***}$ | $0.311^{***}$ | $0.404^{***}$ |
| MPD | $0.109^{***}$ | $0.325^{***}$ | $0.667^{***}$ | $0.843^{***}$ |
| **Fungi** | | | | |
| Richness | $0.099^{***}$ | $0.200^{***}$ | $0.445^{***}$ | $0.579^{***}$ |
| MPD | $-0.064^{*}$ | $-0.115^{*}$ | $-0.305^{***}$ | $-0.586^{***}$ |
| **Considering abiotic variables** (Fit: $\alpha$-MF ~ $\alpha$-diversity + abiotic variables)**:** | | | | |
| | $\alpha$-MF (avg.) | $\alpha$-MF (25%) | $\alpha$-MF (50%) | $\alpha$-MF (75%) |
| **Bacteria** | | | | |
| Richness | $0.043^{*}$ | n.s. | $0.570^{***}$ | $0.751^{***}$ |
| MPD | n.s. | n.s. | n.s. | n.s. |
| **Cyanobacteria** | | | | |
| Richness | n.s. | $0.138^{**}$ | n.s. | n.s. |
| MPD | $0.026^{*}$ | $0.193^{*}$ | $0.398^{***}$ | $0.466^{**}$ |
| **Fungi** | | | | |
| Richness | $0.031^{**}$ | $0.143^{**}$ | $0.302^{***}$ | $0.397^{**}$ |
| MPD | n.s. | n.s. | n.s. | n.s. |

The significant levels are indicated as $^{***}$ $p < 0.001$, $^{**}$ $p < 0.01$, $^{*}$ $p < 0.05$.

**Supplementary Table 5.** Comparison between multiple models for microbial *β*-diversity when selecting geographic distance (Geodist) or ΔMAP for analysis. *D*s: total *β*-diversity; *Repl*: species replacement; $R_{diff.}$: richness difference; ΔMAP: variance in mean annual precipitation; ΔFreq: variance in mean annual precipitation frequency; ΔVar: variance in precipitation variability.

| **Formula** | adj $R^2$ | *p*-values | AICs |
|---|---|---|---|
| **Bacteria** | | | |
| *D*s ~ ΔElevation+ΔMAP+ΔVar+ΔFreq+ΔpH+ΔSalinity+ΔC:N | 0.313 | < 0.001 | 2448.51 |
| *D*s ~ Geodist+ΔElevation+ΔVar+ΔFreq+ΔpH+ΔSalinity+ΔC:N | 0.572 | < 0.001 | 1979.22 |
| *Repl* ~ ΔElevation+ΔMAP+ΔVar+ΔFreq+ΔpH+ΔSalinity+ΔC:N | 0.367 | < 0.001 | 2366.46 |
| *Repl* ~ Geodist+ΔElevation+ΔVar+ΔFreq+ΔpH+ΔSalinity+ΔC:N | 0.548 | < 0.001 | 2033.53 |
| $R_{diff}$ ~ ΔElevation+ΔMAP+ΔVar+ΔFreq+ΔpH+ΔSalinity+ΔC:N | 0.318 | < 0.001 | 2440.03 |
| $R_{diff}$ ~ Geodist+ΔElevation+ΔVar+ΔFreq+ΔpH+ΔSalinity+ΔC:N | 0.315 | < 0.001 | 2444.88 |
| **Cyanobacteria** | | | |
| *D*s ~ ΔElevation+ΔMAP+ΔVar+ΔFreq+ΔpH+ΔSalinity+ΔC:N | 0.233 | < 0.001 | 2557.27 |
| *D*s ~ Geodist+ΔElevation+ΔVar+ΔFreq+ΔpH+ΔSalinity+ΔC:N | 0.372 | < 0.001 | 2358.48 |
| *Repl* ~ ΔElevation+ΔMAP+ΔVar+ΔFreq+ΔpH+ΔSalinity+ΔC:N | 0.210 | <0.001 | 2585.59 |
| *Repl* ~ Geodist+ΔElevation+ΔVar+ΔFreq+ΔpH+ΔSalinity+ΔC:N | 0.330 | < 0.001 | 2422.71 |
| $R_{diff}$ ~ ΔElevation+ΔMAP+ΔVar+ΔFreq+ΔpH+ΔSalinity+ΔC:N | 0.126 | < 0.001 | 2685.61 |
| $R_{diff}$ ~ Geodist+ΔElevation+ΔVar+ΔFreq+ΔpH+ΔSalinity+ΔC:N | 0.126 | < 0.001 | 2686.02 |
| **Fungi** | | | |
| *D*s ~ ΔElevation+ΔMAP+ΔVar+ΔFreq+ΔpH+ΔSalinity+ΔC:N | 0.303 | < 0.001 | 2461.79 |
| *D*s ~ Geodist+ΔElevation+ΔVar+ΔFreq+ΔpH+ΔSalinity+ΔC:N | 0.550 | < 0.001 | 2027.87 |
| *Repl* ~ ΔElevation+ΔMAP+ΔVar+ΔFreq+ΔpH+ΔSalinity+ΔC:N | 0.198 | < 0.001 | 2601.56 |
| *Repl* ~ Geodist+ΔElevation+ΔVar+ΔFreq+ΔpH+ΔSalinity+ΔC:N | 0.323 | < 0.001 | 2433.17 |
| $R_{diff}$ ~ ΔElevation+ΔMAP+ΔVar+ΔFreq+ΔpH+ΔSalinity+ΔC:N | 0.060 | < 0.001 | 2758.35 |
| $R_{diff}$ ~ Geodist+ΔElevation+ΔVar+ΔFreq+ΔpH+ΔSalinity+ΔC:N | 0.065 | < 0.001 | 2752.65 |

**Supplementary Table 6.** Effects of abiotic and biotic (community composition) variables on distance-based *β*-multifunctionality (Euclidean) based on multiple regression. ΔMAP: variance in mean annual precipitation; ΔFreq: variance in annual precipitation frequency; ΔVar: variance in daily precipitation variability; *Repl*: species replacement; $R_{diff}$: richness difference.

| Predictors | Bacteria | Cyanobacteria | Fungi |
|---|---|---|---|
| **Standard coefficients of predictors:** | | | |
| ΔElevation | 0.008 | -0.019 | -0.038 |
| ΔMAP | -0.134$^{***}$ | -0.120$^{***}$ | -0.076$^{***}$ |
| ΔVar | 0.243$^{***}$ | 0.341$^{***}$ | 0.236$^{***}$ |
| ΔFreq | -0.104$^{***}$ | -0.141$^{***}$ | -0.070$^{**}$ |
| ΔpH | 0.125$^{***}$ | 0.147$^{***}$ | 0.144$^{***}$ |
| ΔSalinity | 0.410$^{***}$ | 0.310$^{***}$ | 0.302$^{***}$ |
| ΔC:N | -0.004 | -0.043$^{*}$ | -0.014 |
| *Repl* | 0.513$^{***}$ | 0.514$^{***}$ | 0.552$^{***}$ |
| $R_{diff}$ | 0.314$^{***}$ | 0.325$^{***}$ | 0.474$^{***}$ |
| $R^2$ | 0.635 | 0.632 | 0.651 |
| **Adjusted** $R^2$ | 0.632 | 0.629 | 0.648 |
| ***F*-statistic** | 189.4 | 187.2 | 202.9 |
| ***p*-value** | < 0.001 | < 0.001 | < 0.001 |

The significant levels are indicated as $^{***}$ $p < 0.001$, $^{**}$ $p < 0.01$, $^{*}$ $p < 0.05$.

**Supplementary Table 7.** Effects of abiotic and biotic (community composition) variables on asynchrony-based $\beta$-multifunctionality ($\beta$-MF) based on GLMs. ΔMAP: variance in mean annual precipitation; ΔFreq: variance in annual precipitation frequency; ΔVar: variance in daily precipitation variability; *Repl*: species replacement; $R_{diff}$: richness difference.

**Standard coefficients of predictors:**

| | Bacteria | | | Cyanobacteria | | | Fungi | | |
|---|---|---|---|---|---|---|---|---|---|
| **Predictors** | $\beta$-MF (25%) | $\beta$-MF (50%) | $\beta$-MF (75%) | $\beta$-MF (25%) | $\beta$-MF (50%) | $\beta$-MF (75%) | $\beta$-MF (25%) | $\beta$-MF (50%) | $\beta$-MF (75%) |
| ΔElevation | 0.05** | -0.02 | 0.02 | 0.02 | -0.04** | 0.01 | 0.03 | -0.05** | 0.00 |
| ΔMAP | 0.01 | -0.06*** | -0.10*** | 0.01 | -0.04** | -0.10*** | 0.06*** | -0.02 | -0.13*** |
| ΔVar | -0.05** | 0.08*** | 0.37*** | 0.00 | 0.15*** | 0.37*** | -0.03 | 0.06** | 0.28*** |
| ΔFreq | 0.07*** | -0.10*** | -0.28*** | 0.04* | -0.13*** | -0.28*** | 0.08*** | -0.06*** | -0.23*** |
| ΔpH | 0.05** | 0.00 | 0.08*** | 0.07*** | 0.03 | 0.09*** | 0.10*** | 0.02 | 0.04* |
| ΔSalinity | 0.06*** | 0.15*** | 0.19*** | 0.00 | 0.08*** | 0.18*** | 0.01 | 0.07*** | 0.17*** |
| ΔC:N | 0.07*** | -0.07*** | -0.10*** | 0.05*** | -0.09*** | -0.10*** | 0.07*** | -0.07*** | -0.11*** |
| $R_{diff}$ | 0.26*** | 0.24*** | -0.01 | 0.28*** | 0.19*** | -0.02 | 0.22*** | 0.36*** | 0.25*** |
| *Repl* | 0.31*** | 0.37*** | 0.09*** | 0.31*** | 0.33*** | 0.13*** | 0.20*** | 0.45*** | 0.34*** |
| Nagelkerke's $R^2$ | 0.515 | 0.664 | 0.723 | 0.520 | 0.630 | 0.735 | 0.394 | 0.701 | 0.761 |

The significant levels are indicated as *** $p < 0.001$, ** $p < 0.01$, * $p < 0.05$.
